# Replaying germinal center evolution on a quantified affinity landscape

**DOI:** 10.1101/2025.06.02.656870

**Authors:** William S. DeWitt, Ashni A. Vora, Tatsuya Araki, Jared G. Galloway, Tanwee Alkutkar, Juliana Bortolatto, Tiago B.R. Castro, Will Dumm, Chris Jennings-Shaffer, Tongqiu Jia, Luka Mesin, Gabriel Ozorowski, Juhee Pae, Duncan K. Ralph, Jesse D. Bloom, Armita Nourmohammad, Yun S. Song, Andrew B. Ward, Tyler N. Starr, Frederick A. Matsen, Gabriel D. Victora

## Abstract

Darwinian evolution of immunoglobulin genes within germinal centers (GCs) underlies the progressive increase in antibody affinity following antigen exposure. Whereas the cellular mechanics of how competition between B cells produces increases in affinity are well established, the evolutionary dynamics of this process are less clear. We developed an experimental evolution model where we “replay” over one hundred monoclonal GC reactions, assigning affinities to each cell using deep mutational scanning. Our data reveal how GCs achieve predictable outcomes by means of noisy but persistent selection on an affinity landscape whose exploration is heavily constrained by somatic hypermutation biases. We infer a fitness landscape that quantitatively recapitulates the affinity maturation trajectory of our clone and find that apparent features of GC selection such as permissiveness to low-affinity lineages and rapid plateauing of affinity are likely artifacts of survivorship biases that distort our view of how B cell affinity progresses over time.

## INTRODUCTION

The immune system generates vast repertoires of immunoglobulins (Igs) by stochastic gene recombination and subsequently improves—or matures—their affinity for antigen by rapid somatic evolution^1,2^. Affinity maturation takes place in germinal centers (GCs), clusters of rapidly dividing B cells that arise in secondary lymphoid organs upon infection or immunization^2–4^. Within GCs, B cells undergo iterative cycles of somatic hypermutation (SHM) of *Ig* genes followed by selective expansion of mutant lineages with improved affinity. Predicting the outcome of this evolutionary process is an important goal for vaccine design^5,6^.

Although we have developed a broad cellular and molecular framework for how GC evolution operates^2–4^, the dynamical evolutionary principles at play in the GC are not completely understood. For example, “clonal bursts”—abrupt proliferations in which the descendants of a single cell take over the GC within a few days—occur in some GCs but not others, in a manner not easily predictable from the affinity of the bursting GC B cell^7^. Likewise, empirical studies consistently show coexistence of high- and very low-affinity GC B cells within the same lymph node or even the same GC^7–11^, a degree of permissiveness that fits poorly with current mechanistic models of GC selection^2^.

GCs also provide a model system for studying evolutionary dynamics more broadly. Inferring the quantitative relationship between phenotype and fitness—i.e. building a “fitness landscape”^12^—is a central challenge in evolutionary biology, as genotype-to-phenotype-to-fitness mapping is confounded by complex trait spaces, pleiotropy, and the diversity of adaptive strategies that arise even in tightly controlled experimental systems^13–19^. In contrast, GC B cells evolve by rapidly mutating only two *Ig* genes—the heavy chain (*Igh*) and light chain (either *Igk* or *Igl*)—with predictable biases^20–23^, and apply selection on a single trait, ability to bind antigen^24^. Multiple GCs form independently within the same animal, evolve affinity by several orders of magnitude over a few weeks, and leave behind evolutionary trajectories that can be reconstructed from *Ig* sequencing and phylogenetic analysis^25^, providing both abundant replicates and a means to track their history. GCs are therefore a tractable system in which to test the predictability of evolution in a physiologically relevant setting.

We performed a “parallel replay” experiment on GC B cells, which we use as a platform for experimental evolution. Despite high variability among GC phylogenies, selection for high affinity occurred consistently across GCs. Rather than being driven by clonal bursts, affinity maturation resulted from imperfect but persistent selection of affinity-increasing mutations. Combining phylogenetic reconstructions with a fitness landscape inferred from populations sampled over time, we show that both the apparent permissiveness of GCs to low-affinity lineages and the apparent early plateau in affinity maturation are best explained by survivorship biases that distort the histories of lineages present at sampling.

## RESULTS

### Parallel replay of evolutionary trajectories in clonally identical GCs

A quantitative analysis of GC selection requires the ability to generate replicated GC evolutionary trajectories starting from the same pool of founder B cells. To achieve this, we established a system in which GCs are composed entirely of B cells carrying the same pre-rearranged *Igh* and *Igk* genes, ensuring identical starting specificity and affinity. We chose a B cell receptor (BCR) specific for the model antigen chicken IgY (clone 2.1)^7,26^. We engineered mice carrying the rearranged, unmutated *Igh* and *Igk* genes of clone 2.1 in their respective loci, which we refer to as “chIgY” mice (**Fig. S1A,B**). We then bred these mice to a ubiquitously expressed photoactivatable (PA)GFP transgene to enable isolation of B cells from individual GCs within a LN by *in situ* photoactivation^7,24^. To generate monoclonal GCs, we adoptively transferred 5 x 10^5^ purified chIgY B cells (*Igh*^chIgY/+^.*Igk*^chIgY/+^.PAGFP-tg) into CD23-Cre.*Bcl6*^flox/flox^ recipients, which lack endogenous GC B cells^27,28^.

We immunized recipients subcutaneously with IgY in alum to generate GCs consisting almost exclusively of donor cells within an otherwise polyclonal host (**Fig. 1A** and **Fig. S1C**). GCs were analyzed at 15 and 20 days post-immunization (dpi)—roughly 5 and 10 days after GCs peak in cell numbers. We explanted draining LNs, photoactivated 1-4 individual GCs per node using labeled follicular dendritic cells (FDCs) for guidance, sliced nodes into segments containing a single photoactivated GC, then sorted photoactivated B cells from each segment into 96-well plates for sequencing (**Fig. 1B** and **Fig. S1C**). We obtained paired *Igh* and *Igk* sequences for 8,744 B cells from 119 replicate GCs (15 dpi: 3,758 cells from 52 GCs (18 mice), median 75 (range 30-87) cells/GC; 20 dpi: 4,986 cells from 67 GCs (6 mice), median 78 (25-94) cells/GC). Somatic mutations were present in most cells (10 and 1 unmutated cells at 15 and 20 dpi, respectively). Median SHM load was 5 (range, 0-18) and 7 (0-19) nucleotide mutations per cell at 15 and 20 dpi, respectively.

**Figure 1.**
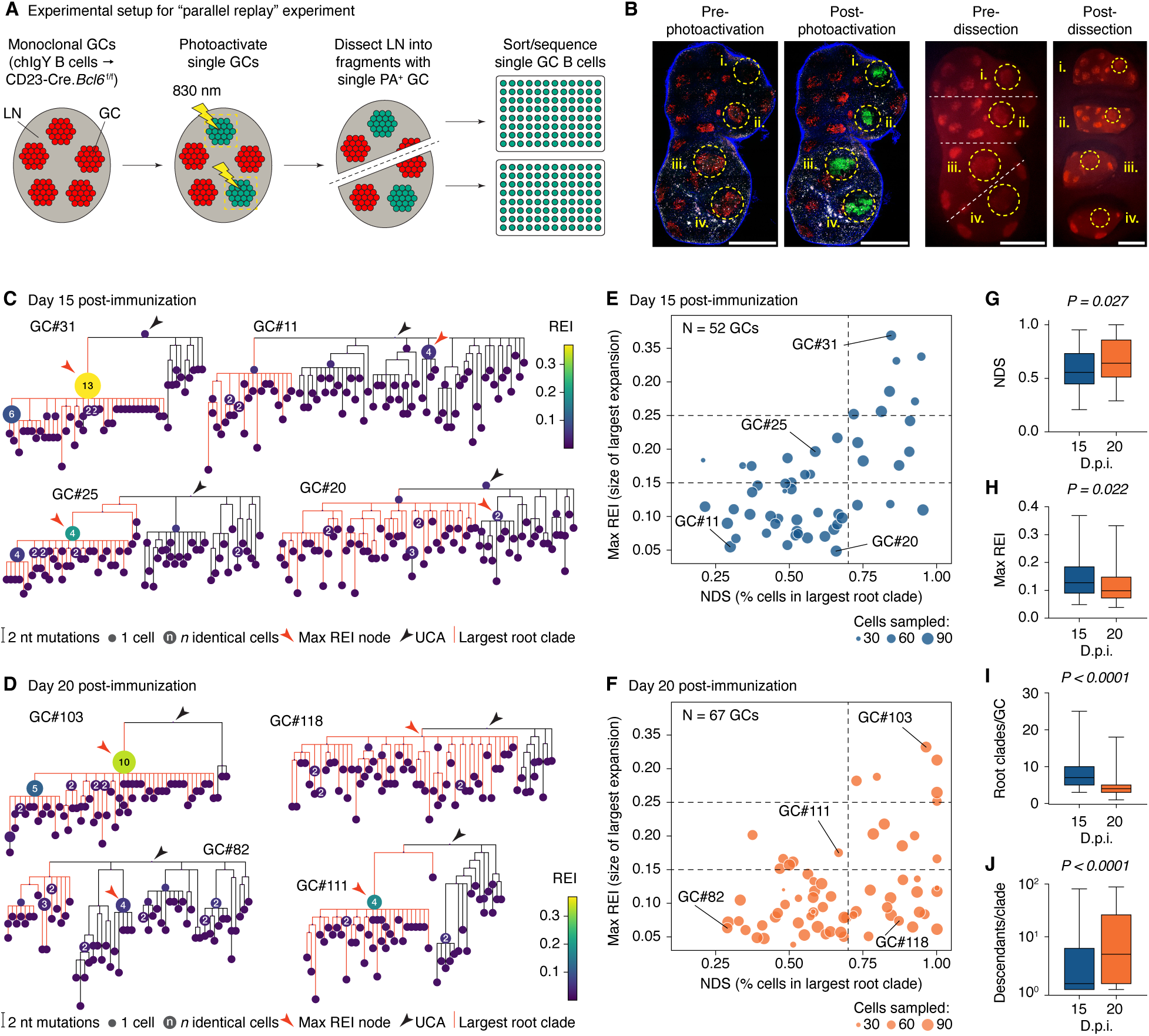
Parallel replay of germinal center evolution. **(A)** Experimental design. Monoclonal germinal centers were generated by transfer of IgY-specific B cells into GC-deficient (CD23-Cre.*Bcl6*^flox/flox^) mice, which are then immunized with IgY to generate GCs. At 20 dpi, one or more individual GCs per LN were photoactivated under a multiphoton microscope. LNs were then dissected into fragments containing a single photoactivated GC, and photoactivated GC B cells were sorted into 96-well plates for *Ig* sequencing. **(B)** Example of GC photoactivation. Left, tiled multiphoton images showing photoactivation of four individual GCs in one node. Image is one Z-plane. Right, fluorescent stereoscope images of LNs prior to and after dissection into fragments. Dotted lines and roman numerals indicate photoactivated GCs. Scale bar, 0.5 mm. **(C,D)** Examples of phylogenetic trees for GCs sequenced at 15 and 20 dpi. **(E,F)** Distribution of NDS and REI scores for GCs from 15 and 20 dpi; each symbol represents one GC scaled according to the number of B cells sequenced. **(G-J)** Comparison of phylogenetic features of GCs obtained at 15 and 20 dpi. Bar is median, boxes are 25-75%, whiskers are range. Units are GCs (G-I), and root clades (J). Data are for 52 GCs (3,758 cells; 413 root clades) from 18 mice, 2 independent experiments for 15 dpi and 67 GCs (4,986 cells; 271 root clades) from 6 mice in 2 independent experiments for 20 dpi. P-values are for Mann-Whitney U test.

To quantify reproducibility across replicates, we first inferred phylogenetic trees for each GC using the *gctree* package^25,29,30^. This revealed a wide variety of topologies, ranging from large clonal bursts to highly branched trees with multiple lineages stemming from the unmutated ancestor (**Fig. 1C,D** and **Fig. S2**). We quantified this variation using two metrics, a “normalized dominance score,” denoting the size of the largest lineage in the GC, and the maximum “recent expansion index” (max REI), which measures the magnitude of the largest burst in that GC (**Fig. S1D,E**). GCs displayed a wide range of NDS and REI scores at both time points (**Fig. 1E,F**), which was consistent between mice and across different LNs (**Fig. S1F**). GCs with low NDS and max REI had multiple, evenly competitive root-clades (e.g., GCs #11, #82), indicating failure of any single clade to establish dominance. At the other extreme were large clonal bursts (e.g., GCs #31, #103), the strongest of which were able to eliminate most competing clades. Although the frequency of clonal bursts (top-right sector, e.g., GC #31) was similar between time points (7/52 at 15 dpi vs. 6/67 at 20 dpi), the fraction of GCs with high NDS but low max REI (bottom-right sector; e.g. GC #118) increased significantly at 20 dpi (3/52 vs. 18/67; p_Fisher_ = 0.0031). This was accompanied by modest decreases in max REI and number of root-clades per GC, alongside slight increases in NDS and number of descendants per root-clade, from 15 to 20 dpi (**Fig. 1G-J**). Thus, as established clonal bursts “age,” their descendants accumulate mutations, reducing max REI. On the other hand, root-clade diversity (equivalent to the diversity of V(D)J rearrangements in a polyclonal setting) is not restored, such that NDS remains high.

We conclude that the evolutionary trajectories of GCs are highly variable even when founder populations are identical at the *Ig* sequence level, clonal bursts being observed at frequencies similar to those previously reported^7,31^. Thus, a simplified monoclonal model recapitulates the range of outcomes observed for polyclonal GCs.

### Determining the effects of somatic mutation on antigen binding

To understand how features of clone 2.1 phylogenies related to changes in affinity, we used deep mutational scanning (DMS) to measure the impact of virtually all possible V(D)J single-amino acid (aa) replacements to the naïve 2.1 sequence on IgY binding and Ig surface expression (**Fig. S3A**). We constructed a mutagenesis library containing 4,158 out of 4,161 possible single-aa replacements available to clone 2.1, cloned into a single-chain fragment variable (scFv) construct for yeast-surface display^32^. We then the Tite-Seq approach^33^ to determine the impact of each replacement on IgY binding affinity (Δaffinity, defined as –Δlog_10_(K_D_)) and surface scFv expression levels (Δexpression, a proxy for folding stability^34,35^) (**Fig. 2A** and **S3A-C**).

**Figure 2.**
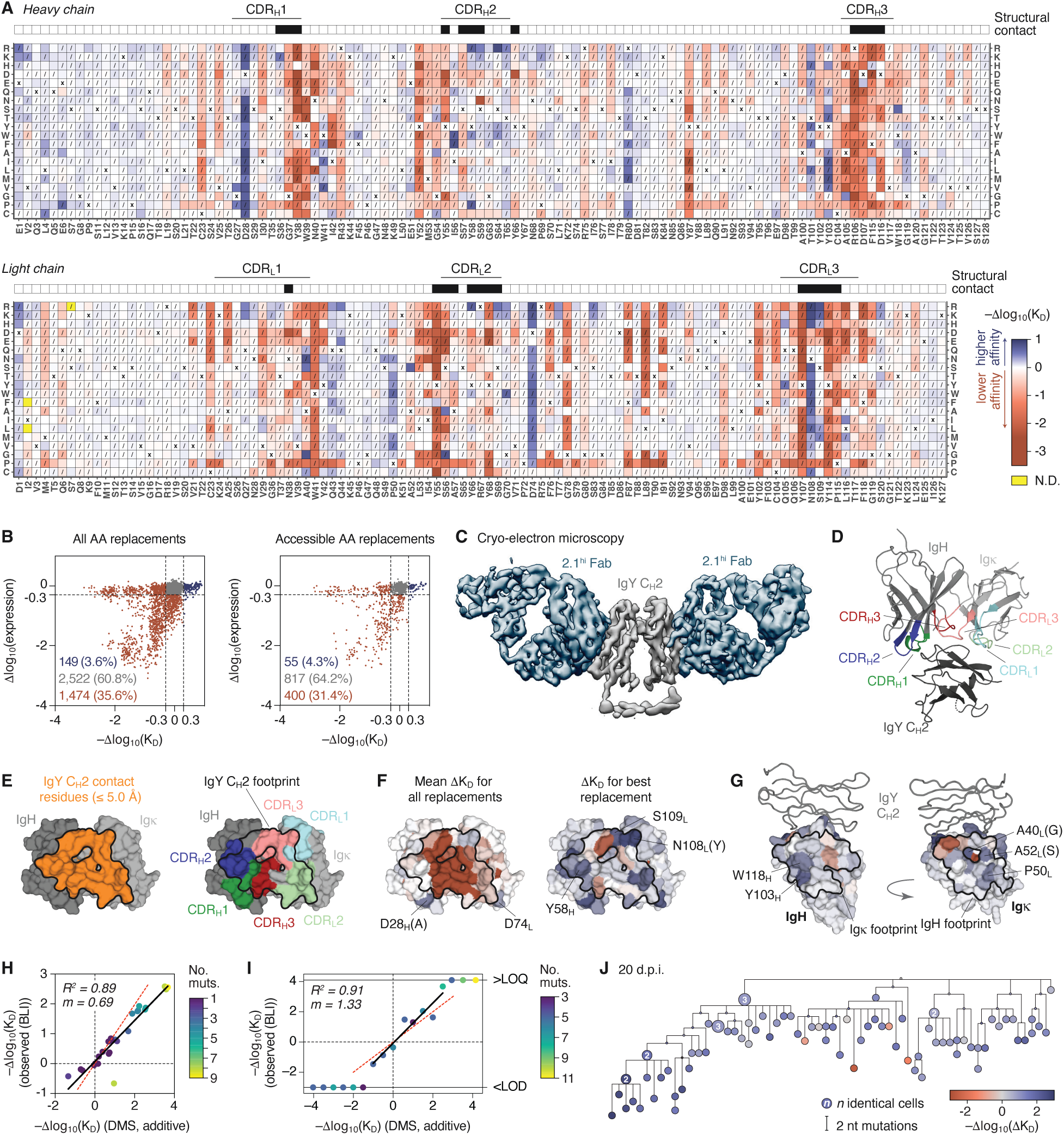
Deep mutational scanning and structure of clone 2.1. **(A)** Heatmaps showing the effects of individual amino-acid replacements on binding of 2.1 scFv to chicken IgY by yeast display. Each square represents a different replacement. Squares marked with “X” indicate the original amino acid in clone 2.1. Squares with slashes indicate amino acid replacements distant more than one nucleotide mutation from the naïve sequence. Yellow squares were not detected in the DMS experiments. Antigen-antibody contact residues (intermolecular distance ≤ 5.0 Å as determined by cryo-EM) are shown as black boxes in the upper bar. See Fig. S2C for the equivalent heatmap for scFv expression. An interactive version of this heatmap is available at https://matsengrp.github.io/gcreplay/interactive-figures/mutation-heatmaps/naive_reversions_first.html; replacements can be visualized on the Fab structure at https://matsen.group/gcreplay-viz/. Data are the mean of two independent experiments. **(B)** Effects of all (left) and accessible (requiring a single nucleotide mutation) amino acid replacements on 2.1 affinity and surface expression. The number and fraction of amino acid replacements in each category (impairment, red; neutral, gray; improvement, blue) are indicated. **(C)** 4.0 Å cryo-EM reconstruction of the 2.1^hi^ Fab (D28HA, K49HR, S64HG, A40LG, Y42LF, A52LS, Q105LH, N108LY) complexed to IgY following local refinement using a mask around the IgY CH2 and 2.1^hi^ Fab regions. **(D)** Cryo-EM structure of the interaction of the 2.1^hi^ V-domain with IgY CH2 with CDRs indicated (PDB: 9ODB). **(E)** Structure of the 2.1^hi^ V-domain showing the footprint (black outline) of CH2, defined as residues with intermolecular distance ≤ 5.0 Å. The antibody-antigen interface spans 851 Å2 (341 Å2 for VH and 510 Å2 for V_k_). **(F)** As in (E), colored by mean Δaffinity (–Δlog10(KD) for all replacements (left), or maximum Δaffinity for the best possible replacement at each position (right)). Color scale as in (A). **(G)** Structure of the 2.1^hi^ V-domain heavy (left) and light (right) chains colored by maximum Δaffinity. The approximate footprint of the opposite antibody chain (residues with ≥ 5 Å^2^ buried surface area contribution) is indicated by a black outline. IgY CH2 is shown as a gray ribbon. Color scale as in (A). **(H-I)** Comparison of Δaffinities predicted using the additive DMS model and BLI measurements of Fabs produced recombinantly using the same sequences. Solid black line is the linear trend; dotted red line is *x = y*. (H) 2.1 variants found frequently in prior studies of clone 2.1; (I) Affinity ladder spanning 8 orders of magnitude. LOD, limit of detection, below which BLI curve fitting was unreliable; LOQ, limit of quantitation, when Fab off-rate is too long to be determined. R^2^ and slope (*m*) calculations exclude Fabs <LOD and >LOQ. **(J)** Example GC phylogeny from the 20 dpi replay dataset (see Fig. 1), colored based on the additive DMS model.

As expected given the relatively high affinity of unmutated clone 2.1 (∼40 nM^7^), aa changes, particularly those falling within complementarity-determining regions (CDRs), were much more likely to reduce affinity than to improve it (**Fig. 2A,B**). Whereas the best available replacement improved affinity by less than one log_10_ (N108_L_R, Δaffinity = 0.91), the worst lowered affinity by 3 log_10_ (Y38_H_E, Δaffinity = –3.0; **Fig. 2A**). Of the 4,145 aa replacements assayed for both Δaffinity and Δexpression in the DMS, 1,474 (35.6%) led to at least a 0.3 log_10_ decrease in either binding affinity or scFv surface expression, while only 149 (3.6%) led to a gain in affinity of 0.3 log_10_ or greater (**Fig. 2B**). Similar results were obtained when only accessible replacements (those resulting from a single nucleotide mutation, non-hatched squares in Fig. 2A) were considered (400 (31.4%) deleterious and 55 (4.3%) enhancing replacements of 1,272 assayed; **Fig. 2B**). Thus, for every enhancing replacement clone 2.1 can make, it must avoid making roughly 10 deleterious ones. No aa replacements were found that led to a substantial gain in surface expression, suggesting that stability of clone 2.1 is near-optimal (**Fig. 2B** and **S3C**).

To understand the structural basis for the DMS results, we determined negative-stain and cryo-EM structures of an affinity-matured version of the 2.1 Fab (measured K_D_ = 62 pM) bound to IgY. Clone 2.1 bound to the hinge-like C_H_2 domain of the four-domain IgY constant region (**Fig. S3D,E**). The paratope of clone 2.1 consisted of a concave pocket that contacted the outer face of IgY C_H_2 (**Fig. 2C-E** and **Fig. S3F**). Mapping the DMS to the structure showed that replacements in the central groove of the paratope had strongly negative mean effects on Δaffinity, whereas affinity-enhancing replacements were found primarily at the periphery of the paratope (**Fig. 2E,F** and **Fig. S3G**) and along the heavy-light chain interface (**Fig. 2G**). Replacements that reduced scFv surface expression occurred in their expected positions (e.g., disulfide-bond cysteines and inward-facing hydrophobic residues and salt bridges^36^; **Fig. S3H**).

To estimate the affinities of GC B cells containing multiple mutations, we simply added the Δaffinities associated with each individual aa replacement, reasoning that, since most affinity-enhancing replacements in clone 2.1 were located along the edges of an otherwise optimal central groove (**Fig. 2F**), epistatic interactions leading to non-additive effects would be limited. To validate this approach, we produced a series of recombinant Fabs carrying selected affinity-enhancing replacements that appear frequently in clone 2.1, either alone or in combination (references ^7,26^ and our unpublished observations) and measured their binding to IgY by biolayer interferometry (BLI **Supplemental Spreadsheet 1**). This showed good agreement (R^2^ = 0.89, slope = 0.69) between predicted and observed values within this series (**Fig. 2H**). We next produced a 17-step “ladder” of Fabs derived from B cell sequences obtained from the replay experiment, spanning Δaffinities between –4.0 and +4.0. Fabs with estimated Δaffinity < –1.0 bound too weakly to antigen to be characterized. Above this threshold, DMS-predicted and BLI-measured affinities increased linearly up to Δaffinity ≅ 2.5 (R^2^ = 0.91, slope = 1.33; **Fig. 2I**), after which off-rates were too long to accurately measure. Overall, the mean absolute difference in Δaffinity between BLI measurements and DMS predictions was 0.17 for antibodies with a single mutation and 0.64 for antibodies with more than one mutation. (As an estimate of the accuracy of BLI, the mean absolute difference between 9 independent BLI measurements of the unmutated Fab and the mean of these measurements was 0.20.) The predicted mean Δaffinity for all antibodies tested was 1.06, compared to a measured mean of 0.88, a difference of 0.18. We conclude that adding the DMS-determined effects of individual mutations is sufficiently accurate to predict how multiple mutations affect the affinity of clone 2.1, particularly when these affinities are averaged across many cells. **Fig. 2J** shows an example of a 20 d.p.i. GC phylogeny, colored using this model (for all trees, see **Fig. S2**).

### Selection of individual aa replacements across germinal centers

Using the DMS, we assessed how efficiently chIgY GCs identified and selected for each of the available affinity-enhancing aa replacements. Because the observed frequency of a replacement in the population depends on both its Δaffinity and on the intrinsic mutability of its codon given SHM targeting biases^37,38^, we first measured nucleotide mutability across each of the *Ig*^chIgY^ alleles in the absence of selection using “passenger” alleles (*Igh*^chIgY^* and *Igk*^chIgY^*) containing frameshifts in the leader sequence upstream of each V-region (**Fig. S4A**). These alleles are transcribed and mutated but do not produce functional proteins, and can thus be used to measure the intrinsic mutability of each nucleotide in an *Ig* sequence in the absence of antigen-driven selection^37^. **Fig. 3A** shows relative mutation rates for each nucleotide in *Igh*^chIgY^* and *Igk*^chIgY^* (obtained by sequencing GC B cells induced by *Plasmodium chabaudi* infection, see Methods). As expected, intrinsic mutability varied greatly across each sequence and was generally higher in CDRs than in frameworks (**Fig. 3A**). Observed mutation rates correlated significantly but not perfectly with those predicted using a five-mer context model^23^ (**Fig. S4B**).

**Figure 3.**
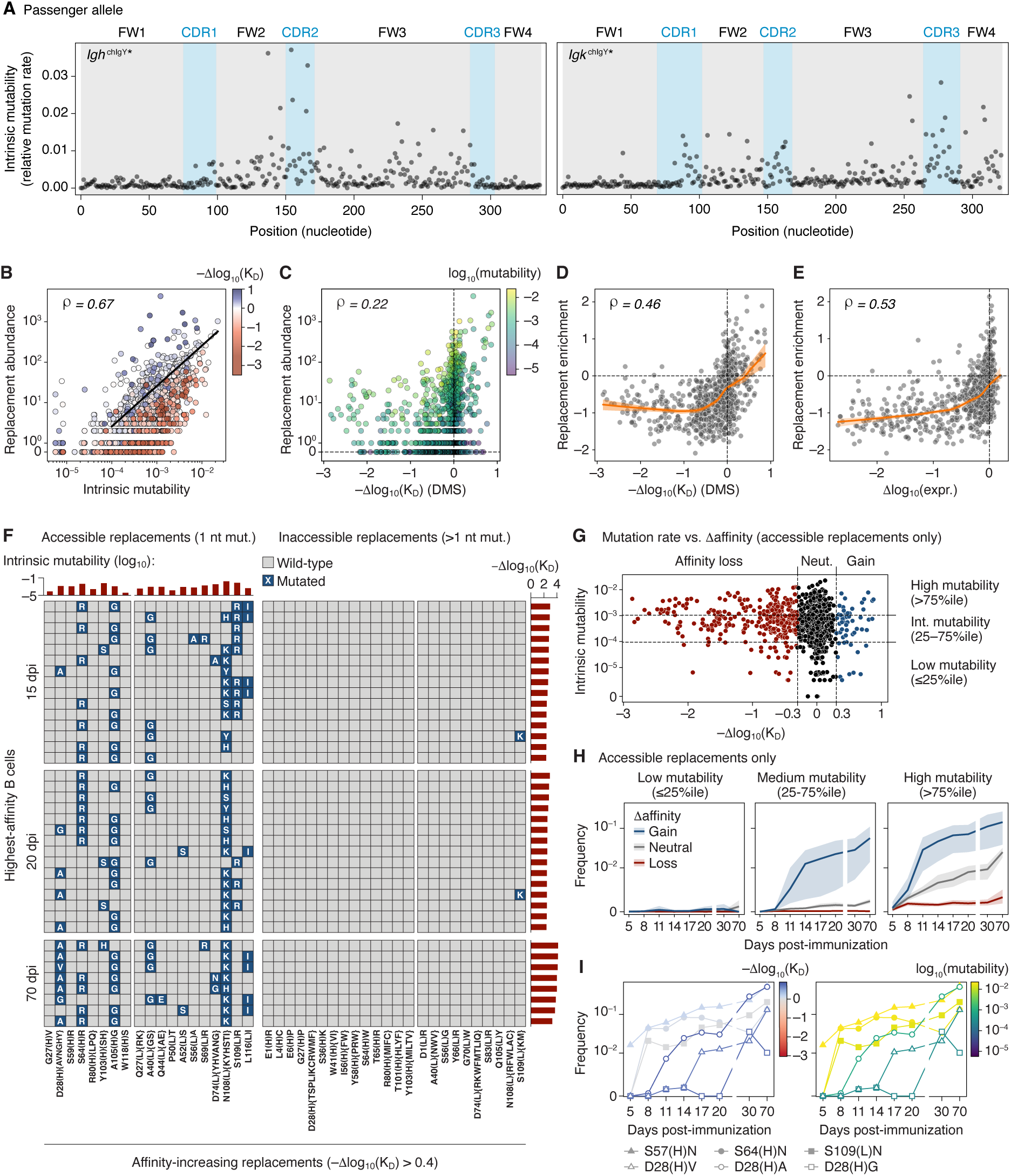
Accumulation of beneficial replacements in germinal centers is constrained by mutability. **(A)** Relative mutation rate measured for each base pair of the passenger *Igh*^chIgY^* and *Igk*^chIgY^* alleles. Each symbol represents one nucleotide position, the sum of the relative rate of mutation for each of the three non-native nucleotides is given. Each symbol represents one nucleotide position, the sum of the relative rate of mutation for each of the three non-native nucleotides is given. Data are pooled from 3 mice for *Igh* and 2 mice for *Igk*. **(B)** Correlation between relative mutation rate, calculated for each of 1,275 codon-accessible amino acid replacements based on the data in (A), and the prevalence of each of these replacements in the replay dataset. Each symbol represents one replacement, colored by the Δaffinity determined for that replacement by DMS. Black line is a Poisson regression with no link function, used to calculate the “replacement enrichment” in (D,E). **(C)** Correlation between Δaffinity and the prevalence of each of these replacements in the replay dataset. Each symbol represents one replacement, colored by relative mutation rate. **(D,E)** Correlation between Δaffinity (D) and Δexpression (E) and replacement enrichment (defined as the log_10_ of the number of observed events over the number of predicted events according to a Poisson regression, shown as a black line in (A)). The orange line shows a LOWESS regression with 95% confidence intervals. ρ values are for Spearman correlation. **(F)** Accumulation of affinity-enhancing replacements among the highest-affinity B cells at various time points post-immunization. Each row represents the Ig sequences of one cell (the highest-affinity B cell in its host GC). Each column represents the set (curly brackets in the X-axis label) of affinity-enhancing replacements (affinity > 0.4 according to the DMS) available at each position, grouped into those accessible by a single nucleotide mutation (left) and those that are not (right). Grey squares indicate that a B cell has the WT amino acid at that position; blue squares indicate replacements, and the identity of the replacement made by the B cell is given in white font. Bars above the graph indicate the summed intrinsic mutability for the listed replacements calculated from the passenger allele. Bars to the right indicate the DMS-estimated Δaffinity of each B cell. **(G)** Classification of accessible amino acid replacements into categories of Δaffinity (DMS) and relative mutation rate (passenger allele), defined by the dashed lines. **(H)** Accumulation over time of replacements from the 9 categories in (F) among GC B cells sequenced at various time points after immunization as detailed in **Figure S4G**. Each line represents the average normalized frequency per mutation category calculated as each mutation’s total frequency divided by the total number of cells captured at each time point. The shaded area represents the 95% CI. Data are pooled from 4 mice per time point and 9 mice for the 70-day time point. **(I)** As in G, but showing accumulation over time of selected individual replacements.

Intrinsic nucleotide mutability was a much stronger predictor of the frequency of aa replacements in GCs *in vivo* than the Δaffinity associated with each replacement (Spearman ρ = 0.67 vs. 0.22, respectively; **Fig. 3B,C**). Nevertheless, replacements that improved affinity (blue in **Fig. 3B**) were clearly enriched above the regression line, whereas deleterious replacements (red) were enriched below. We therefore plotted the observed frequency of a replacement against its predicted frequency based on intrinsic mutability, defining “replacement enrichment” as the log_10_ fold-change over the regression line. Replacement enrichment was better correlated with Δaffinity (Spearman ρ = 0.46) than observed frequency alone (**Fig. 3C,D**) and also correlated well with Δexpression (Spearman ρ = 0.53; **Fig. 3E**). Both correlations were distinctly biphasic, starting out relatively flat but steepening markedly as they approached zero (**Fig. 3D,E**). Presence of such “breakpoints” was confirmed using segmented regression (**Fig. S4C**). In the range surrounding neutrality, a 10-fold change in affinity or expression led to on average 15- and 100-fold (1.2 and 2.0 log_10_) changes in replacement enrichment, respectively. Thus, when replacements are analyzed in aggregate, GCs appear responsive to even small changes around neutral phenotypic values, but counterselection is near maximal already at moderate expression or affinity loss. Of note, although affinity and expression were strongly correlated for many replacements, within a given range of affinity loss, replacements that also caused loss of expression were more likely to be counterselected (**Fig. S4D**).

To determine how efficiently GCs identified and selected for the full complement of affinity-enhancing aa replacements available to them, we collected the 15 highest-affinity B cells from each replay time point (allowing only one cell per GC) and analyzed their acquisition of the full set of 104 replacements associated with affinity gains of at least 0.4 log_10_ (**Fig. 3F**). GCs largely failed to find affinity-enhancing aa replacements that required more than a single nucleotide mutation in the same codon. Only one of 64 such replacements (S109_L_K, located within the highly mutable CDR_L_3 region) was observed in this sample (**Fig. 3F**), and more generally, only 156 (1.8%) of 8,744 B cells sequenced in the replay experiment carried any of these 64 replacements. Low intrinsic mutability also prevented GCs from finding several of the beneficial replacements accessible with single-nucleotide mutations. These included positions G27, R80, and W118 in IgH and Q27 and P50 in Igk (**Fig. 3F**). To investigate whether these replacements might be selected for if given enough time, we immunized mice as in the replay experiment but using chicken IgY in alhydrogel adjuvant, which generates longer-lived GC responses^39^, then sequenced GC B cells pooled from entire LNs 10 weeks later. These cells were heavily mutated (median 18.5 nucleotide mutations, range 11-37) and some had very high predicted affinities (Δaffinity >4.0), indicative of prolonged GC selection. Nevertheless, no inaccessible or low-mutability aa replacements were detected in the highest-affinity B cells from each 70-dpi sample; rather, these cells achieved high affinities primarily through combinations of mutations also found at earlier time points.

To explore these trends systematically, we generated an independent dataset consisting of a time-course of chIgY GC B cells obtained from mice immunized as in the replay experiment but sorted in bulk (i.e. B cells from multiple GCs were pooled from whole LNs of several mice per time point). Sorted cells were analyzed by droplet-based single-cell sequencing at 5, 8, 11, 14, 17, 20, 30 and 70 dpi, the last two time points using alhydrogel as an adjuvant to extend GC lifetime (**Fig. S4E,F**). We then used passenger allele and DMS data to categorize all accessible aa replacements based on mutability (low (<25%ile), medium (25-75%ile), and high (>75%ile)) and Δaffinity (negative (<–0.3), neutral (–0.3 to 0.3), positive (>0.3)), respectively (**Fig. 3G**). In line with our previous analysis, low-mutability replacements were largely ignored by GCs, even when leading to substantial gains in affinity (**Fig. 3H**). By contrast, enrichment for affinity-enhancing replacements was evident in the high and intermediate mutability classes. High-mutability replacements accumulated progressively over time even when neutral (but not when deleterious), again indicating strict counterselection of affinity-reducing replacements. Following selected replacements over time revealed a pattern where neutral but high-mutability replacements (such as the S to N changes in positions S57_H_, S64_H_ and S109_L_) accumulated early on, whereas unlikely but beneficial ones (such as D28_H_ to V, A, or G) caught up only much later (**Fig. 3I**).

In summary, affinity maturation is heavily constrained by intrinsic biases in SHM and accessibility through single-nucleotide changes. These constraints limit the exploration of the full mutational landscape, even under prolonged selection.

### Phylogenetic analysis identifies the drivers of affinity maturation

#### GCs select consistently for increases in affinity and maintenance of Ig expression

Whereas GC phylogenies varied widely in structure (**Fig. 1D,F** and **Fig. S2**), replay GCs were much more consistent with respect to affinity maturation. Median affinity was higher than naïve in 117 of 119 GCs, while variance was relatively low (0.87 ± 0.38 at 15 dpi and 1.00 ± 0.34 at 20 dpi (mean ±SD); **Figs. 4A** and **S5A**). This consistency was more evident when observed trees were displayed alongside neutral drift simulations in mutational load vs. Δaffinity “trajectory plots” (**Fig. 4B**). In these simulations, replacements are introduced according to mutability alone, in the absence of affinity-based selection, while maintaining the phylogenetic structure of each GC. Neutral simulations consistently produced median affinities markedly lower than observed experimentally (Δaffinity = –0.47 ± 0.24 at 15 dpi and –0.70 ± 0.26 (mean ±SD); **Fig. 4C**). Thus, despite the strong downward pressure on affinity exerted by stochastic mutagenesis, GCs consistently achieved increases in affinity over time.

**Figure 4.**
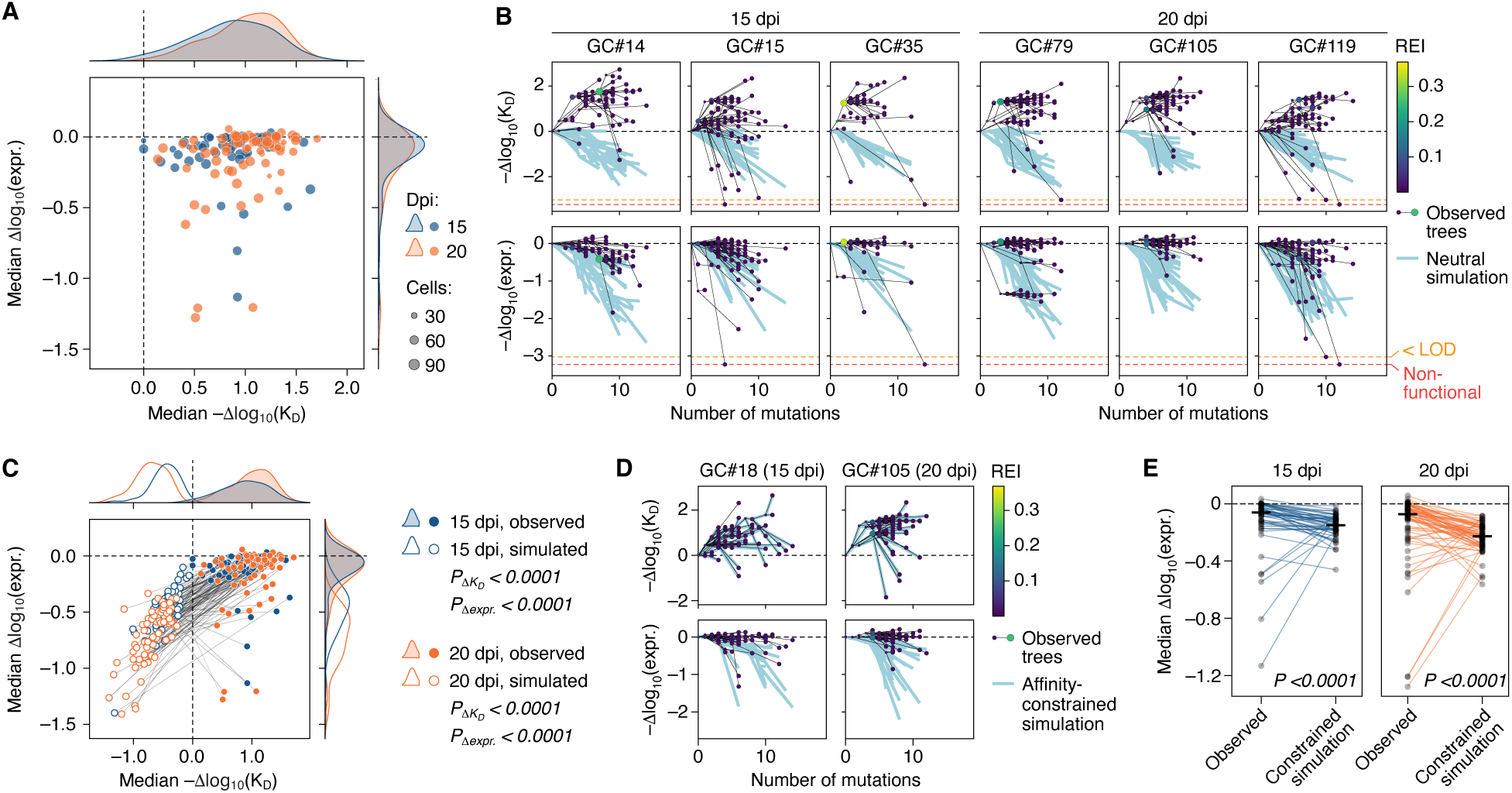
Quantifying germinal center selection for affinity and for maintenance of Ig expression. **(A)** Distribution of 119 replicated GCs by median affinity and Ig expression, as quantified using the additive DMS model. Each symbol represents one GC, symbol sizes are proportional to the number of cells sequenced. Densities on the top and right show the distribution of GCs according to Δaffinity and Δexpression, respectively. **(B)** Example trajectory plots in which phylogenetic trees are plotted in 2D space according to their number of somatic mutations and Δaffinity (top) or Δexpression (bottom). For the experimentally observed trees, circles representing individual nodes, colored by REI and scaled by number of identical sequences, are connected by black lines. Simulated trees with the same phylogenetic structure but in which mutations are assigned based on mutability alone are shown as light blue lines, which represent the median values for each node in 10 simulated trees. Plots for all GCs available at https://github.com/matsengrp/gcreplay/tree/main/results/notebooks/phenotype-trajectories/naive_reversions_first. **(C)** Distribution of 119 replicated GCs by median affinity and Ig expression as in (A) (filled symbols), paired to the medians of 10 simulated GCs as in (B) (open symbols). P-values are for the Wilcoxon signed-rank test. **(D)** Example trajectories as in (B), but simulations are constrained by affinity—i.e., replacements are assigned based on nucleotide mutability but must match the Δaffinity of the observed node within 0.05 log10(ΔKD). **(E)** Comparison of median Δexpression between experimental GCs and affinity-constrained simulations as in (D). Each symbol represents one experimental GC (“observed”) or the median of 10 simulated GCs (“simulated”).

SHM led to a noticeable decrease in Ig surface expression, consistent with previous observations^40^. Median Δexpression dropped below naïve in 106 of 119 GCs (–0.13 ± 0.21 at 15 dpi and –0.17 ± 0.27 at 20 dpi (mean ±SD)), with several instances of more substantial loss (**Fig. 4A-C** and **S5A**). In the four GCs in which median Δexpression fell below –1.0 (**Fig. 4A** and **S5B**) the median-expression B cell acquired replacement Y42_L_N (arrowhead in **Fig. S4D**), which causes a severe drop in expression (–1.27) while increasing antigen binding (0.26). Thus, even strongly destabilizing replacements can still undergo positive selection if they improve affinity. Δexpression was also much lower in simulated than in experimental trees (–0.48 ± 0.22 at 15 dpi and –0.73 ± 0.25 at 20 dpi (mean ±SD); **Fig. 4C**). Because affinity and expression are partially correlated (**Fig. 2B**), we performed additional simulations in which we forced simulated trees to match the Δaffinity of experimental GC lineages within 0.05 log_10_(K_D_), regardless of effects on Δexpression. Median Δexpression in these simulations remained marginally higher than *in vivo* (medians, – 0.060 vs. –0.15, at 15 dpi and –0.073 vs. –0.23 at 20 dpi, p < 0.0001 for both; **Fig. 4D,E**), indicating some degree of selection acting specifically on Ig expression.

In conclusion, despite wide variation in tree shapes, GCs reliably select for increased antibody affinity, despite the downward pressure of random mutation. GCs also exert some independent pressure to maintain Ig expression, though large losses in expression can be tolerated if compensated by affinity gains, consistent with previous work^40^.

#### Low-affinity B cell lineages are efficiently terminated

A recurring feature of GC trajectory plots are steep downward-trending lines leading to terminal nodes lacking detectable descendants (for example, GCs #15 and #119 in **Fig. 4B**), appearing in affinity-colored phylogenies as red-tinted “leaves” at terminal branch ends, downstream of higher-affinity internal nodes (**Fig. 5A** and **Fig. S2**). This pattern suggests that GC lineages that lose affinity are efficiently terminated, rarely leaving detectable descendants. To quantify this, we categorized nodes by Δaffinity relative to their immediate ancestor (which approximates the change in affinity incurred by a B cell in its most recent round of SHM), divided into three categories (<–1.0, –1.0 to 0.3, and >0.3) and plotted the mean mutational distance to the nearest branch tip per category. At both time points, nodes that lost affinity compared to their immediate ancestors were disproportionately enriched at leaves (i.e., more likely to be at distance zero from the closest branch tip, and thus to have been recently generated; **Fig. 5B**). We conclude that B cells that lose affinity due to SHM rarely persist in GCs. By extension, the low-affinity B cells frequently observed in our dataset likely represent recent mutants not yet purged from the population by selection.

**Figure 5.**
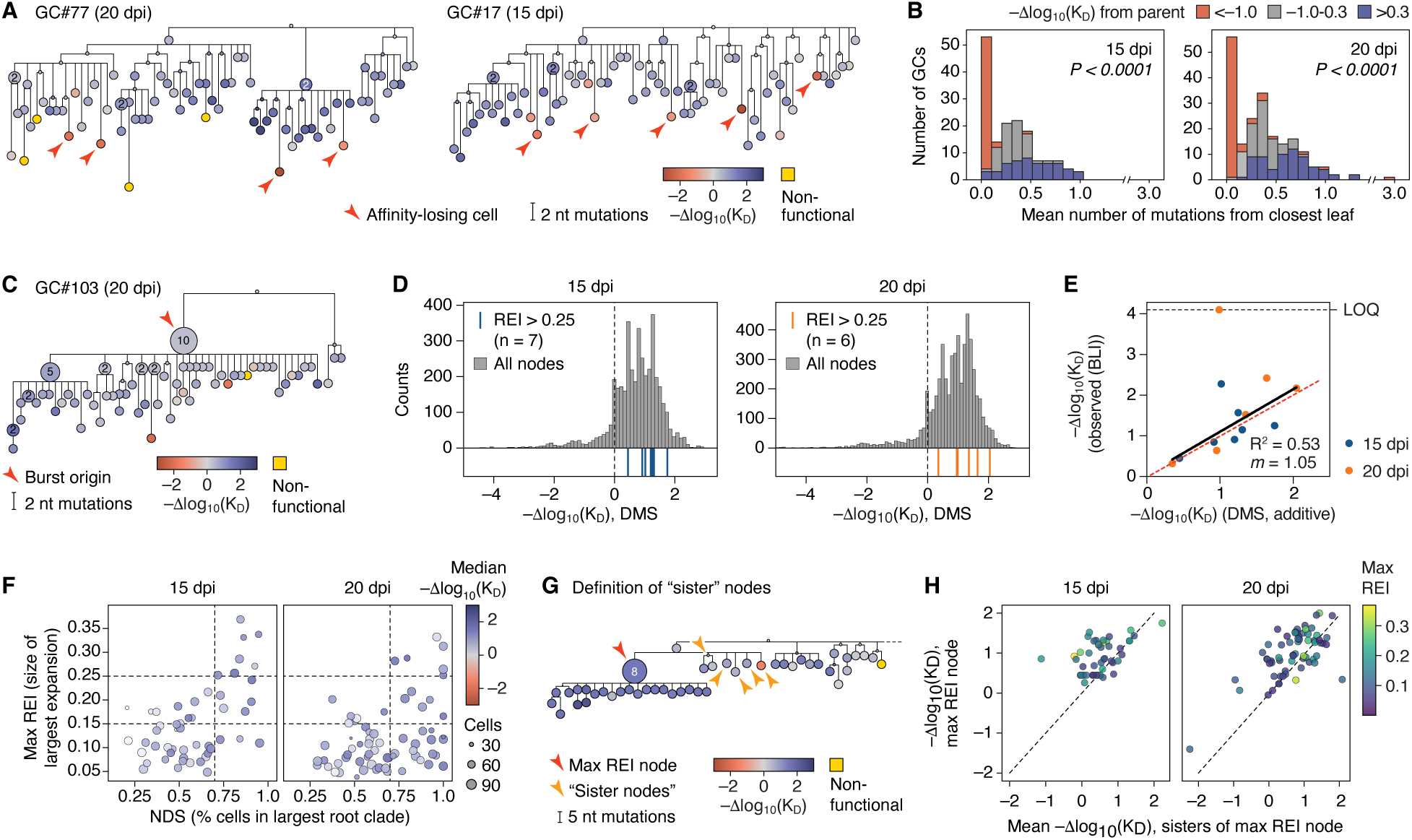
Drivers of affinity maturation as determined by phylogenetic analysis. **(A)** Example phylogenies colored by Δaffinity, indicating cells that lost substantial affinity and are located at terminal nodes (red arrowheads). **(B)** Observed and inferred nodes from each replay GC phylogeny were divided into those that lost (<–1.0), maintained, (–1.0 to 0.3) or gained (>0.3) affinity with respect to their parent node. For each GC, the mean distance between the nodes in each category and the nearest leaf was then recorded (a distance of 0 indicates that the node is itself a leaf). The plot shows the distribution of mean values for each category in each GC. **(C)** Example of a clonal burst phylogeny colored by Δaffinity. The burst point is indicated by a red arrowhead. **(D)** Distribution of Δaffinities for clonal burst nodes (REI >0.25; blue lines for 15 dpi and orange lines for 20 dpi) compared to the distribution of Δaffinities for all nodes (observed and inferred) from the same time point (grey bars). **(E)** Comparison of Δaffinities predicted using the additive DMS model and BLI measurements of Fabs produced recombinantly based on the *Ig* sequences of each of the bursting nodes in (D). Solid black line is the linear trend; dotted red line is *x = y*. LOQ, limit of quantitation, when Fab off-rate is too long to be determined. R^2^ and slope (*m*) calculations exclude the Fab with Δaffinity >LOQ. **(F)** Distribution of NDS and max REI scores for GCs from 15 and 20 dpi as in Fig. 1E and F but colored by median Δaffinity; each symbol represents one GC scaled according to the number of B cells sequenced. **(G)** Schematic explaining the definition of “sister” nodes used in (H) and Fig. S5F. **(H)** Distribution of replay GCs according to the Δaffinity of the max-REI node in the GC and the mean Δaffinity of its “sister” nodes, as defined in (G). For a statistical comparison see Fig. S5F.

#### Clonal bursts are not the primary drivers of affinity maturation

Clonal bursts represent extreme “jackpot” events in which a single B cell proliferates enough to eliminate most competing lineages in its GC^7^,^41^. If burst size were reliably determined by affinity, sporadic large bursts could drive affinity maturation by increasing mean population affinity in a stepwise manner. Counter to this notion, the largest clonal bursts at 15 and 20 dpi (**Fig. 1E,F**, top-right sector) originated from B cells with relatively high but far from extreme affinities (mean percentile rank 66 (range 30–94) at 15 dpi and 64 (21–97) at 20 dpi; **Fig 5C,D**). Bursts from 20 dpi were also not outliers when compared to the 15-dpi background (**Fig. S5D**), indicating that bursting cells were also unexceptional at the time they were selected^42^. BLI-measured affinities of recombinant Fabs derived from 13 clonal bursts (**Fig. 5E)** were relatively well predicted by the DMS, (R^2^ = 0.53, slope = 1.05) with the exception of one Fab whose affinity fell outside the range of the instrument. There also was no correlation between max REI and BLI affinity (**Fig. S5E**). Moreover, color-coding GCs in the NDS x max REI scatter plot (**Fig. 1E,F**) by median Δaffinity revealed no marked enrichment for high-affinity cells in the upper right “clonal burst” sector (**Fig. 5F**). Similarly, there was no strong correlation between affinity measures and burst size (REI) (**Fig. S5C**). Thus, clonal bursts are not a superior strategy for affinity maturation compared to more gradual evolution. Of note, Δaffinity correlated moderately with NDS, especially at 15 dpi (**Fig. S5C**), indicating that evolving multiple lineages simultaneously is associated with lesser affinity gains. We conclude that clonal bursts are neither primary drivers of affinity maturation nor inherently superior as a strategy to less punctuated evolution.

Given the absence of a strong link between clonal bursting and affinity gain, we sought to identify common drivers of affinity maturation present in all GCs, regardless of large-scale tree topology. To this end, we selected the highest-REI B cell in each GC as well as all of its “sister” nodes within the same branch (nodes that shared a common immediate ancestor with the node of interest), which we reasoned would be representative of the competitors of the highest-REI B cell at the time it was selected (**Fig. 5G**). We then measured the gain in affinity between each node and its sisters and that of their immediate ancestor. This showed that, although the discrimination between bursts and sisters was imperfect (i.e., the highest-REI node did not always have higher affinity than its sisters), it was on aggregate significantly skewed towards expanding higher-affinity B cells over their lower-affinity competitors (**Fig. 5H** and **S5F**).

Taken together, our data indicate that affinity maturation in GCs, rather than being driven by sporadic clonal bursts, results from persistent selection that is relatively inaccurate but sufficiently biased towards high affinity B cells to reliably favor their expansion over time.

### A fitness landscape for the temporal evolution of germinal center affinity

The distribution of B cell affinities in a GC evolves over time due to three dynamical factors: *Ig* sequence mutations by SHM, the affinity effects of these mutations, and the fitness effects of affinity differences. This last factor—the “fitness landscape”^12^—is a key conceptual tool in evolutionary dynamics but is rarely directly inferred^43,44^. We sought to infer this landscape for clone 2.1 by assigning affinities to all cells in the time-course experiment (**Fig. S4E,F**), which sampled pooled GC B cells from whole LNs at multiple time points (**Fig. 6A**, gray histograms and **Fig. S6A**). We formulated a minimal mathematical model^45–48^ that predicts the continuous distribution of affinity *x* (–Δlog_10_(K_D_) from naïve) among GC B cells over time *t* (dpi). We denote this distribution 𝑝(𝑥, 𝑡) and formulate a partial differential equation for its temporal evolution, incorporating: (i) a mutation process specified by mutation targeting biases from the passenger allele experiment and the affinity effects from the DMS experiment **(Fig. 6B**, upper panel); and (ii) an affinity-fitness landscape 𝑓(𝑥) specifying fitness for each affinity 𝑥 (**Fig. 6B**, lower panel). The growth (or decay) rate of a B cell population with affinity 𝑥 at time 𝑡 is 𝑓(𝑥) − 𝑓̅(𝑡), where 𝑓̅(𝑡) is the mean fitness of the full population at time 𝑡 (**Fig. S6C**). We used maximum likelihood to estimate 𝑓(𝑥) so that the predicted 𝑝(𝑥, 𝑡) matches our time-course data (**Fig. 6A**).

**Figure 6.**
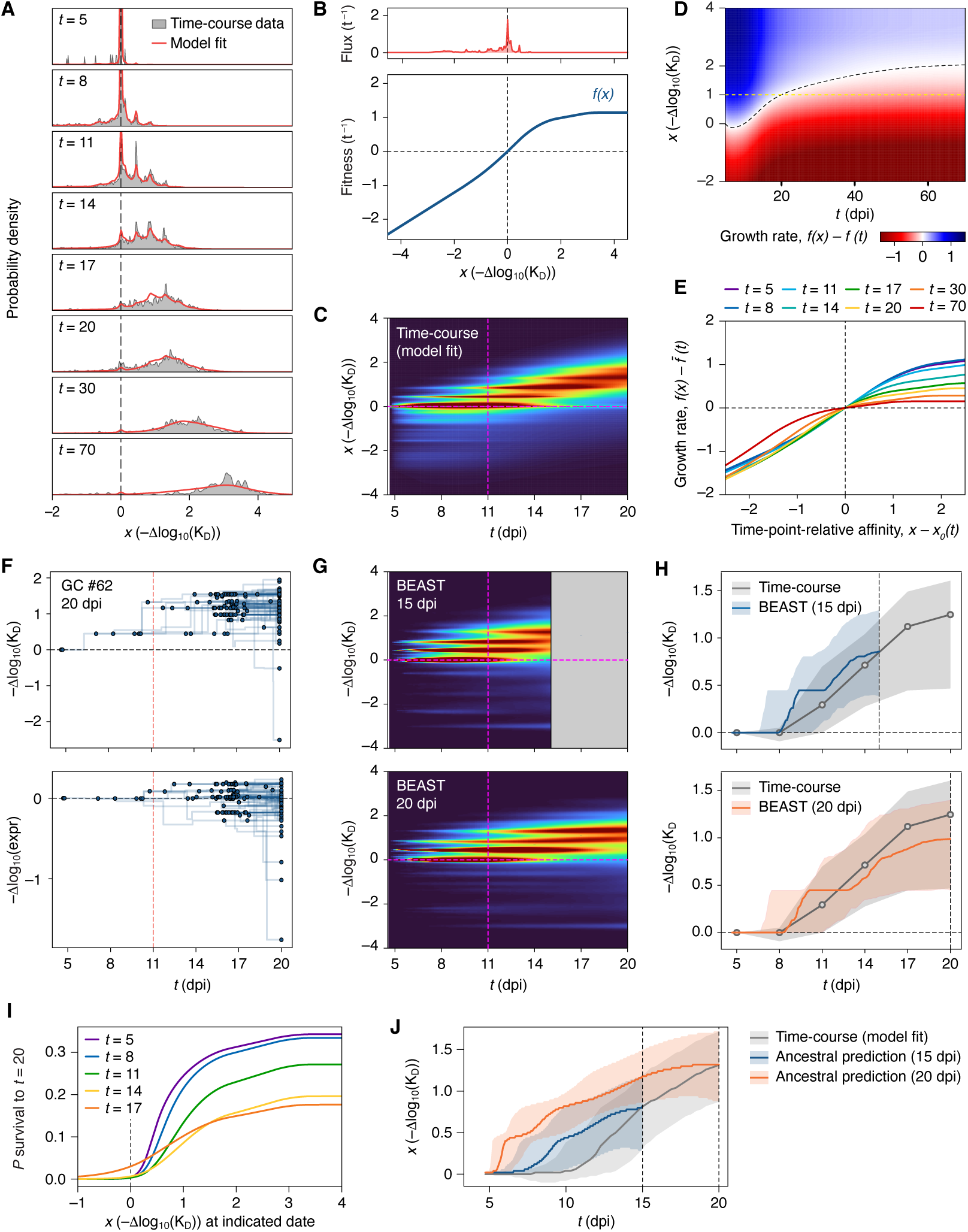
An affinity–fitness landscape inferred from the evolution of clone 2.1. **(A)** Distributions of affinity at 5, 8, 11, 14, 17, and 20 dpi in the time-course experiment (grey; see Fig. S4E,F for a description of the experiment) compared to time-slice fits using the fitness landscape model (red). **(B)** Key parameters driving the fitness landscape model: the distribution of affinity mutations specified by DMS effects and mutation propensities (upper panel), and the fitness landscape (lower panel). **(C)** Solution of the fitness landscape model showing the affinity distribution evolving in continuous time up to day 20 for comparison with (B) and (C). Dotted lines given for reference. **(D)** The growth rate, given as the intrinsic fitness minus the population mean fitness at each time, is shown in the colormap over time and affinity. The affinity corresponding to population mean fitness at each time (i.e., is tracked in the dashed line (we call this the *mean-matched affinity*, and the difference from that, the *time-point-relative affinity*). **(E)** The growth rate, given as the intrinsic fitness minus the population mean fitness, is plotted against the time-point-relative affinity at sampling times. At later times, the fitness response to affinity gains is diminished. **(F)** Example time-resolved tree for one GC showing nodes (birth events) and mutations (affinity jumps) in time (dpi), with all leaves at the sampling time (20 dpi for this GC). 100 such candidate trees were sampled for each GC. Taking a time slice from one such tree (e.g., at the red dotted line) yields a collection of ancestral cells and their associated affinities. **(G)** Heatmaps showing the evolution of affinities over time resulting from aggregating all candidate time-resolved trees for 15 and 20 dpi GCs. Dotted lines given for reference. The push of the past and pull of the present are seen as excess densities in the upper-left and lower-right quadrants, respectively, compared to (F). **(H)** Grey trendline shows median and interquartile range (IQR) affinity in the time-course experiment at 5, 8, 11, 14, 17, and 20 dpi. The blue and orange trend lines show median and IQR affinity through time derived from time-resolved trees for 15 dpi and 20 dpi GCs, respectively. **(I)** Probabilities of survival until 20 dpi for cells of different affinity at various previous times, given the fitted fitness landscape model. These curves are interpreted as the distortions that elevate or suppress different affinities in the reconstructed process. **(J)** The solution of the fitness landscape model (grey) summarized as median and IQR affinity evolving in continuous time. Orange and blue show predicted median and IQR affinity for ancestral population histories sampled at 15 dpi and 20 dpi, respectively, by reweighting the solution with survival probabilities as shown in (G).

The affinity–fitness landscape of clone 2.1 (**Fig. 6B**, lower panel) begins linearly with a slope of 0.74—i.e., a B cell lineage with a 10-fold affinity advantage over competitors will double in size roughly once a day (*e*⁰·⁷⁴ = 2.09-fold/day)—before saturating at around *x* = 2 (400 pM) and reaching a plateau at *x* = 3 (40 pM). Inferring *f(x)* allows us to predict how the affinity distribution would evolve under continuous sampling (**Fig. 6C**) and how the growth potential of B cells changes with time (**Fig. 6D**). The plateauing of *f(x)* at higher affinities implies that the relationship between a B cell’s affinity relative to its competitors (the “time-point-relative affinity”) and its expected growth rate flattens at later time points as the mean affinity enters the pM range (**Fig. 6E**). Thus, selection of improved mutants becomes progressively more difficult later in the GC reaction.

### Phylodynamic survivorship bias distorts tree-based signals of GC B cell fitness

The affinity–fitness landscape allows us to assess how closely time courses reconstructed from phylogenetic trees correspond to those obtained from sequential sampling. We focused on a prominent feature of the GC trajectory plots: many phylogenies appeared to show rapid early gains in affinity followed by plateauing and eventual decline towards the leaves (**Fig. 4B**), suggesting slowdown in GC selection after only a few somatic mutations. To measure these trends over the ensemble of all trees, we first used Bayesian phylogenetics^49^ to generate time-resolved phylogenies for each GC. These time-resolved trees (exemplified in **Fig. 6F**) represent branch lengths as inferred time intervals, placing mutations and nodes at concrete time points in the history of the GC. Because time-resolved tree details have considerable statistical uncertainty, we used 100 posterior tree samples (each a candidate time-resolved tree) for each GC. We used these trees to reconstruct the evolution of B cell affinity over time across all GCs in the replay experiment (**Fig. 6G,H**). This analysis confirmed the observations made from individual trees: affinity-increasing mutations appear to emerge rapidly and are maintained for much of the GC reaction as affinity plateaus (upper-left quadrant in **Fig. 6G**), followed by a subpopulation of lower-affinity B cells at sampling time (lower-right quadrant). Tree-based reconstruction of GC dynamics seems to suggest a rapid change in the strength of affinity-based selection early in the reaction. However, evolution of affinity in this reconstruction differed markedly from that inferred based on the affinity–fitness model (**Fig. 6C**), which decreased slightly in the first days after GC formation—reflecting initial acquisition of deleterious mutations by SHM—followed by gradual increase through day 20. Likewise, we observed the late emergence of cells with negative Δaffinity in the reconstructions (**Fig. 6G**, lower-right quadrant) that were absent from days 14-20 of the fitted time-course (**Fig. S6A,B**). Thus, phylodynamic reconstruction based on an extant population fails to predict the true progression GC B cell affinities over time.

We hypothesized that this inconsistency reflects two concepts from the macroevolution literature: the “push-of-the-past” and the “pull-of-the-present”^50^. Phylogenetic reconstruction inevitably represents a biased history of the subpopulation destined to become ancestors of the cells sampled at the endpoint—this ancestral subpopulation tends to be higher fitness than its contemporaries because it is conditioned on leaving descendants in the present (push-of-the-past). Conversely, the apparent affinity plateau at late time points would result from the presence of low affinity B cells not yet purged from the population (pull-of-the-present). We used the fitness landscape model to predict the effects of both distortions in our replay data. Using a forward master equation approach, in which we consider the fate of a single cell’s descendants competing in the evolving population 𝑝(𝑥, 𝑡), we computed the probability of stochastic extinction of a lineage prior to sampling time at 15 or 20 dpi (**Fig. 6I**). These calculations predicted strong survivorship effects: lineages with 𝑥 < 0 before 17 dpi are very likely to be extinct by day 20, whereas the highest-affinity lineages are most favored for survival at earlier times, when they are more distant outliers from the mean. Conversely, cells with affinities around 𝑥 = 0 at 17 dpi are much more likely to survive to day 20, indicating a lag between acquiring a deleterious mutation and being eliminated from the GC. These calculated survival probability profiles can be used to reweight the time-course prediction 𝑝(𝑥, 𝑡), giving the predicted time evolution 𝑝-(𝑥, 𝑡) of the ancestral population (the subpopulation destined to leave descendants at the later sampling time, 15 or 20 dpi), shown in **Fig. 6J** alongside the full population 𝑝(𝑥, 𝑡) from which they were computed. These predicted ancestral trajectories show clear distortions—early jumps of affinity followed by subsequent slowing down before the sampling time—that are consistent with our findings from inferred trees (**Fig. 6G**).

We conclude that both sampling of current affinities and phylogenetic reconstruction of past trajectories from extant GC B cell populations are subject to survivorship biases that distort our view of selection dynamics. Modeling these effects allows us to separate true biological processes from observational artifacts, providing insight into how mutation and selection shape affinity maturation.

## DISCUSSION

We present the results of an experimental evolution system in which we replay a simplified, monoclonal GC reaction over one hundred times. We find that, even in this simplified setting, GC selection yields widely divergent tree topologies, from clonal burst-type structures to multi-pronged GCs where multiple lineages evolve in parallel. Thus, the diversity of GC outcomes observed in Brainbow GCs^7^ cannot be attributed exclusively to differences in founder populations in a polyclonal setting. By contrast, DMS-based affinity estimation showed that phenotypic evolution across GCs is broadly consistent: virtually all GCs gained affinity compared to the naïve ancestor, despite navigating a landscape that overwhelmingly favors deleterious replacements. Thus, GCs are reproducible with respect to phenotype but not to phylogeny.

A simple additive model in which the log_10_ DMS-measured effects of single replacements are summed, ignoring epistatic interactions, predicted the affinities of clone 2.1 variants carrying multiple replacements with a degree of accuracy sufficient to resolve large-scale features of GC selection. Similar additivity was observed in a recent study of SARS-CoV-2 antibodies that found epistatic effects to be infrequent^51^, but more substantive epistasis has been reported for broadly neutralizing antibodies (bnAbs) against influenza^52,53^. A possible reason for this discrepancy is that most affinity-enhancing replacements in clone 2.1 are distributed around an otherwise optimal paratope (**Fig. 2F**), making them less likely to interact epistatically. Importantly, the degree of epistasis in a given antibody lineage will affect only the accuracy of DMS-based affinity predictions, not the general rules by which GCs select for high-affinity mutants. The evolutionary dynamics we describe for clone 2.1 should therefore hold even in more epistatic settings.

Access to the full universe of affinity-enhancing replacements available to clone 2.1 revealed that GCs “miss” the large majority of replacements with affinity-enhancing potential, primarily due to codon constraints and the low intrinsic mutability of many *Ig* positions due to AID and SHM targeting biases^37,38^. Inaccessibility along these lines has previously been shown to limit the evolution of certain bnAbs to HIV^54,55^. Our data allow us to quantify the full extent of these biases. These findings also suggest that DMS-guided mutagenesis could increase monoclonal antibody affinities well beyond what is obtainable by affinity maturation *in vivo*.

Previous studies have reported high frequencies of low-affinity B cells within GCs, suggesting that GCs may be permissive to low-affinity clones and variants, which would both help lineages traverse local low-affinity “valleys” as well as maintain broader clonal diversity^7–10,56–58^. Our trajectory plots show instead that most low-affinity GC B cells are evolutionary dead-ends, likely representing recent mutants not yet purged by selection. The presence of low-affinity lineages in GCs in previous work may be explained by specificity for cryptic “dark” epitopes^8,59^ generated by antigen degradation or by the blocking of immunodominant epitopes by circulating antibody^60–63^, either of which would decouple a B cell’s *in vitro*-measured affinity from its capacity to retrieve antigen *in vivo*.

Clonal bursts originated from B cells spanning the full spectrum of affinities present ain the GC, and phylogenies containing large bursts did not have higher median affinities than those lacking them. A scenario in which affinity maturation results from sporadic large-scale expansion of very high-affinity clones is therefore inconsistent with our observations. Notwithstanding, these findings do not negate the importance of clonal bursts to GC biology, as they lead to important losses in clonal diversity in polyclonal settings^7,31^ and may also serve as critical sources of GC-derived plasma cells^56^. By contrast, comparing the most successful node of each GC to its sisters revealed a distinct if imperfect association between affinity and number of progeny. Thus, while selection at the level of individual B cell fate decisions may appear noisy^7,41^, this same process reliably drives long-term affinity maturation when repeated across thousands of individual decisions. The multi-step process by which competition for T cell help drives GC selection^24,64^—including the stochastic nature of B cell access to antigen and T cell help in a highly dynamic environment^65–68^—offers ample opportunity for stochasticity to influence the outcomes of competition while preserving the overall selective bias. Thus, the GC operates as an “imperfect cell sorter,” in which individually error-prone fate decisions, when repeated across thousands of B cells, produce reliable long-term affinity gains.

From the standpoint of evolutionary biology, we established an experimental platform able to quantitatively map genotype to phenotype to fitness. This landscape shows, for example, that a 10-fold affinity advantage translates to approximately a doubling of the size of a population each day, with fitness saturating in the high-to-mid-pM range—broadly matching a previously reported affinity ceiling^69,70^. Whether this reflects true saturation or a decline in affinity discrimination over time, as antigen availability declines and T cell help wanes^71^, remains to be resolved. Finally, combining fitness-based modeling with time-resolved reconstruction of ancestral phylogenies revealed systematic survivorship biases, initially described in the context of macroevolution^50^, which obscure the true dynamics of selection. Modeling these biases explicitly allows us to correct for such distortions and better understand how mutation and selection interact to produce reliable affinity maturation.

### Limitations of the study

We trace the outcomes of GC selection acting on one particular B cell clone. The affinity of clone 2.1 for IgY is relatively high (40 nM) but representative of the naïve affinities of clones that dominate secondary responses in mice^72^. The 2.1 affinity–fitness landscape plateaus at a level compatible with prior estimates, suggesting some degree of generalizability of our results to other systems. Nevertheless, it is possible that particular features of GC evolution, such as the association between affinity and clonal bursting and the strictness of counterselection against affinity-losing mutants, differ in settings where mean affinity is lower. Moreover, to ensure strict reproducibility at the clonal level, our GCs are engineered to lack polyclonal competitors. Thus, our setup assays *intra*clonal, but not *inter*clonal, GC competition, ignoring potential factors such as antibody-mediated feedback^60,61^. Further experimentation with different antigen-antibody pairs and using more clonally complex systems will help clarify each of these points.

## ACKNOWLEDGMENTS

We thank C. Ferreira for mouse work, R. de Carvalho for *Plasmodium* infections, D. Rich, J. Gao, and M. Johnson for computational work support, K. Gordon and J.-P. Truman for cell sorting, and the Rockefeller University Comparative Biosciences and Genomics Resource Centers. We thank M. Busslinger and J. Chaudhury for CD23-Cre and A. Dent for *Bcl6*^flox^ mice. Funded by NIH grants R01AI180451, R01AI139117, R01AI119006 (G.D.V.), R01AI146028 (F.A.M.), DP2AI177890 (T.N.S.), R01HG013117 (Y.S.S.), and R35GM142795 (A.N.); and NSF CAREER award 2045054 (A.N.). The Victora laboratory is supported by the Robertson Foundation and the Stavros-Niarchos Foundation Institute for Global Infectious Disease Research at The Rockefeller University. Computing infrastructure at Fred Hutch is funded by ORIP grant S10OD028685. This work benefited from the 2023 “Statistical Physics and Adaptive Immunity” workshop at the Aspen Center for Physics (NSF grant PHY-2210452), and the 2024 “Interactions and Co-evolution between Viruses and Immune Systems” program at the Kavli Institute for Theoretical Physics (NSF grants PHY-2309135 and Gordon and Betty Moore Foundation Grant 2919.02). W.S.D., T.B.R.C., and J.P. were supported by postdoctoral fellowships from the James S. McDonnel, Life Sciences, and Damon Runyon Cancer Research Foundations, respectively. T.N.S. is a Searle Scholar. G.D.V., F.A.M., and J.D.B. are HHMI investigators.

## AUTHOR CONTRIBUTIONS

W.S.D., T.Ara., F.A.M., and G.D.V. conceptualized the study. *In vivo* experiments were led by T.Ara. and A.A.V., with assistance from J.B., L.M., and J.P.. Quantitative analyses were led by W.S.D., F.A.M., G.D.V., and A.A.V., with assistance from J.G.G., T.B.R.C., W.D., C.J.-S., T.J., and D.K.R.. T.N.S. performed the DMS under supervision of J.D.B.. T.Alk. and G.O. performed electron microscopy under supervision of A.B.W.. A.N. contributed to fitness landscape modeling. W.S.D., A.A.V., F.A.M., T.N.S., and G.D.V. wrote the manuscript with input from all authors.

## RESOURCE AVAILABILITY

All unique/stable reagents generated in this study are available from the lead contact upon reasonable request. Jupyter notebooks can be found at https://matsen.group/gcreplay/key-files/#manuscript-figures. Cryo-EM maps and atomic coordinates of Fab 2.1 in complex with IgY are deposited at Electron Microscopy Data Bank (EMDB) and Protein Data Bank (PDB) under accession codes EMD-70353 and 9ODB.

## COMPETING INTERESTS

G.D.V. and J.D.B. are advisors and hold stock of Vaccine Company Inc. J.D.B. consults or has recently consulted for Apriori Bio, Pfizer, GSK, and Invivyd on topics related to viruses, vaccines and viral evolution. J.D.B, and T.N.S. are inventors on Fred Hutch licensed patents (62/935,954 and 62/692,398) related to viral DMS. T.Ara. is an employee of Pfizer Inc.

## METHODS

### Laboratory Methods

#### Mice

*Igh*^2.1^ and *Igk*^2.1^ mice were generated by CRISPR-Cas9 gene-editing in fertilized mouse oocytes. For *Igh*^2.1^, we inserted the pre-rearranged unmutated V_H_ sequence of clone 2.1, preceded by the 443 bp proximal promoter of *Ighv9-4*^73^ in place of the 4 endogenous *Igh* J segments, using two flanking sgRNAs and a single-stranded DNA (ssDNA) megamer (IDT) as described in the easi-CRISPR method^74^. *Igk*^2.1^ mice were generated by inserting the pre-rearranged unmutated V_κ_ sequence of clone 2.1, preceded by the 188 bp proximal promoter of *Igkv3.12*^73^, in place of the 5 endogenous *Igk* J segments, using two sgRNAs and an ssDNA template generated by PCR amplification of a cloned template plasmid followed by opposite-strand digestion using the Guide-it Long ssDNA production system v2 (Takara Bio). Passenger allele mice (*Igh*^2.1^* and *Igk*^2.1^*) were generated using single sgRNAs to introduce indels into the leader sequence of fertilized *Igh*^2.1/+^ and *Igk*^2.1/+^ oocytes, respectively. Sequences of the sgRNA protospacers and ssDNA donor templates used for mouse generation are provided in **Supplemental Spreadsheet 2**. CD23-Cre (Fcer2a-Cre) BAC transgenic mice^27^ (Jax strain #028197) were provided by M. Busslinger (IMP Vienna). *Bcl6*^f/f^ mice^28^ (Jax strain #023727) were provided by A. Dent (U. Indiana). PAGFP-transgenic mice^24^ (Jax strain #022486) were generated and maintained in our laboratory. All animal experiments were approved by the Rockefeller University’s Institutional Animal Care and Use Committee (IACUC). Mice of both sexes were used for all *in vivo* studies without distinction, with the exception that male B cells were never transferred into female recipients to avoid immune-mediated rejection.

#### Cell transfers, immunizations, and Plasmodium infection

Resting splenic B cells were purified by filtering splenocytes through a 70 μm mesh into PBS supplemented with 0.5% BSA and 1mM EDTA (PBE). CD43 MACS beads were used to purify resting B cells from single-cell suspensions according to the manufacturer’s protocol (Miltenyi Biotec). The percentage of *Igh*^2.1/+^/*Igk*^2.1/+^ resting B cells was determined prior to transfer by staining with chicken IgY-BV421 (conjugated in-house) followed by flow cytometry. For all experiments, 5 x 10^5^ *Igh*^2.1/+^/*Igk*^2.1/+^ resting B cells were transferred into each recipient mouse intravenously. 24 hours following cell transfer, mice were immunized subcutaneously in the hind footpads, inner thighs, and/or forearms with 5, 10, and 20 µg, respectively, of chicken IgY (Exalpha Biologicals) precipitated in 1/3 volume of Imject Alum (ThermoFisher Scientific Cat# 77161) to generate similar-sized GCs in popliteal, inguinal, brachial, and axillary nodes. The immunization schemes were the same for the day-70 experiments, except that Imject Alum was replaced with Alhydrogel (InvivoGen) used according to the manufacturer’s protocol. For analysis of SHM in passenger alleles, either *Igh*^2.1^*^/+^*.Igk*^+/+^ or *Igh*^+/+^.*Igh*^2.1^*^/+^ mice were injected intraperitoneally with 10^5^ *Plasmodium chabaudi*-infected red blood cells (BEI Resources Cat# MRA-741).

#### Imaging and Photoactivation

Multiphoton imaging and photoactivation were performed as described previously^7,24^, using an Olympus FV1000 upright microscope fitted with a 25X 1.05NA Plan water-immersion objective and a Mai-Tai DeepSee Ti-Sapphire laser (Spectraphysics). One day prior to photoactivation, PAGFP-transgenic mice were injected intravenously with 10 μg of a non-blocking antibody to CD35 (clone 8C12, produced in house) conjugated to Cy3 to label networks of follicular dendritic cells (FDCs)^75^. A Leica M165FC fluorescence stereomicroscope with a dsRed filter was used to register the location of FDCs. Clusters of CD35-expressing cells were then identified using multiphoton imaging at λ = 950 nm, at which photoactivation does not take place, and three-dimensional regions of interest were photoactivated by higher-power scanning at λ = 830 nm. Lymph nodes were then sliced manually under a Leica M165FC fluorescence stereomicroscope using double-edged safety razor blades (Astra Superior Platinum) for subsequent flow cytometry and sorting.

#### Flow cytometry and cell sorting

For replay time points, sliced LN fragments (15 and 20 dpi) or whole LNs (70 dpi) were placed into microcentrifuge tubes containing 100 μl PBE, macerated using disposable micropestles, and dissociated into single-cell suspensions by gentle vortexing. We then added 100 μl of 2× antibody stain (B220-BV785, TCRβ-APC Cy7, CD38-APC, Fas-PE-Cy7, biotinylated chicken IgY, Streptavidin-BV421, mouse Igk-PE, supplemented with Fc block) to the cell suspension, which was incubated on ice for 30 min. Photoactivated transferred GC B cells were index-sorted into 96-well plates containing 5 μl TCL buffer (Qiagen) supplemented with 1% β-mercaptoethanol. For passenger allele sequencing, spleens from *P. chabaudi*-infected mice were harvested at 20-21 days post-infection. Single-splenocyte suspensions were obtained by forcing spleens through a 70 µm mesh followed by hypotonic lysis of red blood cells using ACK buffer. CD19 magnetic (MACS) beads (Miltenyi Biotec) were used according to the manufacturer’s protocol to enrich for B cells prior to sorting for splenic GC B cells as described above. For 10X Genomics single-cell *Ig* sequencing, LN cell suspensions were stained with individual hashtag oligonucleotide (HTO)-labeled antibodies to CD45 and MHC-I for sample barcoding prior to GC B cell sorting as above. A combination of two HTO’s per sample was used to accommodate for the number of samples in the experiment. Cells were sorted into microfuge tubes with PBS supplemented with 0.4% BSA and were counted for viability by trypan blue staining prior to loading onto a 10X Genomics Chromium Controller.

#### Single-cell PCR amplification and sequencing of Igh^2.1^ and Igk^2.1^ alleles

Sorted single cells were processed and analyzed essentially as described previously^7^, except that specific forward primers were used to amplify chIgY BCR, and both chains were amplified simultaneously for all wells. Primers were limited to the leader sequences of each *Ig*^chIgY^ allele to enable full sequencing of framework (FR)1 regions. Specific primers used were 5’-AGCGACGGGAGTTCACAGACTGCAACCGGTGTACATTCC-3’ (*Igh*^2.1^ V_H_ leader forward recoded) and 5’-AGCGACGGGAGTTCACAGGTATACATGTTGCTGTGGTTGTCTG-3’ (*Igk*^2.1^ Vk leader forward). The first 18 nucleotides of the *Igh*^2.1^ V_H_ leader forward sequence were prepended to the Vk primer so that same barcoding primers could be used for both chains in the next step. After PCR reactions as described previously^7^, pooled PCR products were purified using SPRI beads (0.7× volume ratio), gel-purified, and sequenced with a 500-cycle Reagent Nano kit v2 for single-cell libraries on the Illumina Miseq platform.

#### Bulk PCR amplification and sequencing of Igh^2.1*^ and Igk^2.1*^ passenger alleles

To sequence passenger alleles from a large enough population of GC B cells, we first generated GCs in *Igh*^chIgY^*^/WT^ and *Igk*^chIgY^*^/WT^ mice by infection with *Plasmodium chabaudi*, which generates large numbers of GC B cells in the spleen^76^. We then bulk-sequenced the *Igh* and *Igk* genes of sorted pools of 7.5 x 10^5^ GC B cells at days 20-21 post-infection, when B cells carried on average 2.2 and 2.6 nucleotide mutations per chain, respectively. 750,000 GC B cells were sorted directly into 750 µl of TRIzol LS (Thermofisher #10296010), bulk RNA was extracted using the manufacturer’s protocol. Quality of RNA was checked using TapeStation D1000, and only samples with RIN > 8 were used for generating BCR libraries. The NEBNext Immune Sequencing kit mouse (#E6330S) was used according to the manufacturer’s instructions, with IgM, IgGa, and IgGb primers used to sequence BCR heavy chains and Igk primers for light chains. Each chain was amplified in separately to ensure even amplification. BCR libraries were sequenced using Illumina NextSeq 2000 flow cell P1, to generate 100 million reads.

#### 10X Genomics single-cell *Ig* sequencing

GC B cells sorted as above were sequenced for gene expression and BCR using the 10x Chromium Next GEM Single Cell 5’ Reagent kit v3 with Feature Barcode technology for Cell surface Protein, according to the manufacturer’s protocol. The resulting library was sequenced on a NovaSeq SP (Illumina) flow cell with a minimum sequencing depth of 30,000 reads per cell. De-multiplexing of the samples based on their unique molecular identifier (UMI) and HTO counts were generated with Cell Ranger v6.0.1, v7.0.1 or v8.0.1 with mm11 reference. BCR libraries were also processed with CellRanger “vdj” with default parameters.

#### Recombinant Fab fragment production and affinity measurements

Fabs were produced in one of two ways. For Fig. 2H, *Igh*^chIgY^ and *Igk*^chIgY^ variable regions were synthesized by Twist Bioscience and directly cloned into a custom human IgG1 Fab expression vector, as described previously^7^. Plasmids were transfected into Freestyle Expi-293 suspension cells (Life Technologies), and monoclonal Fab fragments were purified using Ni-NTA beads (GE Healthcare), according to the manufacturer’s protocol. Protein purity was assessed by SDS-PAGE and functional protein concentrations were calibrated using biolayer interferometry on an Octet Red96 instrument using anti-Fab coated sensors (FortéBio). For Figs. 2I and 5E, Fabs were cloned and produced by GenScript based on *Igh*^chIgY^ and *Igk*^chIgY^ variable region sequences and were purified using Capture Select CH1-XL Magnetic Agarose Beads (ThermoScientific). Purity and functional concentrations were measured as mentioned above. BLI affinity measurements were performed as described previously using Octet SAX biosensors^7^. K_D_ values were calculated using a kinetic 1:1 model. Only Fabs with good global fits (R^2^ > 0.98) over multiple concentrations were used for K_d_ calculations. For Fabs that could not fit globally due to their low affinity (n = 3), partial fits were averaged over multiple concentrations and used instead.

#### Yeast-surface display deep mutational scanning

The clone 2.1 scFv was ordered as a yeast codon-optimized gene and cloned into the pETcon yeast surface-display expression vector containing a previously described barcode sequencing landing pad^77^. A site-saturation mutagenesis library was synthesized by Twist Bioscience, precisely encoding every possible aa substitution at each of the sites in the heavy- and light-chain variable domains. In duplicate, N16 barcodes were appended to mutagenesis products and cloned into the vector as previously described^77,78^. The duplicate libraries were electroporated into *E. coli* (NEB C3020K) and plated into a bottlenecked target of 88,000 cfu per library, aiming for an average of 20 redundant barcodes representing each of the possible aa substitutions in the library. PacBio sequencing was used to link N16 barcode to scFv genotype as previously described^77,78^.

Mutation effects on CGG-binding affinity and scFv surface expression were determined by FACS and sequencing as previously described^77,78^. Briefly, libraries were cloned into yeast (*Saccharomyces cerevisiae* strain AWY101^79^), induced for surface expression and labeled with a FITC-conjugated anti c-Myc antibody (Immunology Consultants Lab, CYMC-45F) to label for scFv surface expression, or anti-c-Myc antibody and biotinylated CGG (Exalpha IgY-B) followed by PE-conjugated streptavidin (Thermo Fisher S866) to label for CGG-binding. CGG incubations were set up across 10-fold ligand concentrations from 10^-6^ to 10^-13^ M, plus a 0 M baseline sample. Library cells were sorted into four bins of Myc-FITC (for expression) or SA-PE (for CGG binding) as described^77,78^, plasmid extracted from outgrown cells from each sort bin, and barcode counts in each FACS bin were determined via 50 bp single end sequencing on an Illumina NextSeq. Illumina sequencing counts were processed into mutant effects on surface expression and CGG-binding affinity as detailed in https://github.com/jbloomlab/Ab-CGGnaive_DMS/blob/main/Titeseq-modeling.ipynb. An interactive version of the DMS and Fab structure is available at https://matsen.group/gcreplay-viz/.

#### Negative-stain electron microscopy (EM)

IgY, either alone or in complex with clone 2.1 Fab, was diluted to ∼0.02 mg/ml in Tris-buffered saline and applied to plasma-cleaned, carbon-coated 400 mesh grids. Grids were stained with 2% (w/v) uranyl formate for 30-60 s and blotted. Imaging was performed on an FEI Tecnai Spirit operating at 120-keV, and micrographs recorded with an FEI Eagle 4k CCD camera. Data were processed using Relion 3.0^80^ or cryoSPARC v3.2^81^.

#### Cryo-EM

IgY was incubated with a threefold molar excess of a Fab fragment of a high-affinity variant of clone 2.1 (2.1^hi^, which includes replacements D28_H_A, K49_H_R, S64_H_G, A40_L_G, Y42_L_F, A52_L_S, Q105_L_H, and N108_L_Y) at room temperature for 5 hours. The final concentration of the complex was diluted to 0.06 – 1.0 mg/ml for vitrification. To aid with sample dispersal on the grid, the complex was mixed with lauryl maltose neopentyl glycol (final concentration of 0.005 mM; Anatrace) and deposited on plasma-cleaned Quantifoil 1.2/1.3 300 mesh grids. A Thermo Fisher Vitrobot Mark IV set to 4°C, 100% humidity, 10 s wait time, and a 3-s blot time was used for the sample vitrification process.

Data were collected using Leginon^82^ over 3 separate imaging sessions on a Thermo Fisher Talos Arctica operating at 200 keV and equipped with a Gatan K2 Summit direct electron detector. Combined movies were aligned and dose weighted using MotionCor2^82^. Aligned frames were imported into cryoSPARC v3.2 and the contrast transfer function (CTF) was estimated using GCTF^83^. Particle picking was done by automated picking using templates created from an initial round of 2D classification, then extracted and subjected to multiple rounds of 2D classification for cleaning. An ab initio volume was generated, followed by 3D classification and the best classes were further refined. To further improve the resolution, the maps were subjected to global and local CTF refinements. A mask was created using UCSF Chimera^84^ and cryoSPARC Volume Tools to cover both clone 2.1 Fabs and the central C_H_2 portion of IgY, and used during local refinement without symmetry. A summary of data collection and processing statistics can be found in **Supplemental Table 1.**

Initial models were generated by fitting coordinates from sAbPred^85^ for the Fab and ColabFold^86^ for the C_H_2 dimer into the cryo-EM map. Several rounds of iterative manual and automated model building and relaxed refinement were performed using Coot 0.9.4^87^, Rosetta Relax^88^ and Phenix real_space_refine^89^. Models were validated using EMRinger^90^ and MolProbity^91^ as part of Phenix software suite. IMGT numbering was applied to the antibody Fab variable light and heavy chains. Final refinement statistics and PDB/EMDB deposition codes can be found in **Supplemental Table 1.** Buried surface area was calculated as the mean for the two monomers in the asymmetrical structure, calculated using PISA interface list (https://www.ebi.ac.uk/pdbe/pisa/).

#### Western blotting

Truncated IgY-GFP fusion constructs were generated by cloning each Ig domain of the IgY heavy chain into the XhoI and EcoRI sites of pEGFP-C1-PRKAA1 (Addgene plasmid #30305). 2 µg of each construct was transfected into HEK293T cells using standard calcium phosphate transfection protocols. 24 hours, cells were harvested and 20 µg of protein extracts were loaded onto SDS-PAGE gels and transferred to nitrocellulose membranes using standard wet transfer protocols. All membrane incubations were performed on an orbital shaker. 5% BSA in Tris buffered saline-Tween 20 (TBST) was used to block the membrane for 1 h at 4°C. Both 0.5 µg/ml of recombinant clone 2.1 (A40G) and a 1:5000 dilution of anti-GFP antibody (Biolegend, cat# 902605) were diluted in TBST with 5% BSA for primary staining overnight at 4°C on an orbital shaker. A 1:2,000 dilution of HRP anti-human IgG or a 1:10,000 dilution of HRP anti-mouse IgG antibodies were diluted in TBST with 5% milk for secondary staining against clone 2.1 and anti-GFP antibody, respectively. After 1 h incubation at room temperature on an orbital shaker, membranes were washed, ECL substrate (Pierce) was added, and membranes were exposed for 300 s for signal detection on an Azure c300 imaging system (Azure Biosystems).

#### Computational and Quantitative Methods

Most plots were generated in Jupyter notebooks, and an index of which plots appear in what notebooks can be found in https://matsen.group/gcreplay/key-files/#manuscript-figures. This pipeline hosted at https://github.com/matsengrp/gcreplay/ and these notebooks form a reproducible artifact describing the analysis.

#### Sequencing data pipeline

BCR sequencing data were processed using a custom Nextflow v24.04.3.5916 pipeline to reconstruct clonal relationships within germinal centers. The pipeline takes, as input, raw MiSeq paired-end sequencing reads, which first trimmed to remove the first three bases using fastx_trimmer^92^, and subsequently combined using pandaseq^93^. The resulting sequences were demultiplexed based on plate and well barcodes using the fastx_toolkit^78^, producing individual files for each 96-well plate. Heavy and light chain sequences were then separated by identifying conserved motifs using cutadapt^94^ with a 20% error allowance. To reduce noise and focus on biologically relevant sequences, each well’s heavy and light chain sequences were collapsed to unique sequences, and low-abundance BCRs (fewer than 5 reads by default) were pruned to generate ranked files.

The filtered BCR light and heavy chain sequences were then merged across all wells, maintaining identifiers of their origins, and subsequently annotated using partis^95^, primarily to identify V(D)J gene segments, and somatic mutations compared to the naive sequence. The annotated BCR’s were then processed to generate comprehensive datasets containing paired heavy and light chain information for each germinal center. With this, the workflow then performs downstream phylogenetic inference with GCtree (described next), and all other downstream analysis on the resulting GC trees.

#### Phylogenetic inference and tree-based analysis

We inferred phylogenies on clonal families using gctree v4.3.0. GCtree uses PHYLIP’s dnapars utility to infer a collection of maximum parsimony (MP) phylogenetic trees on observed sequences. GCtree then uses a data structure called a history sDAG to expand the collection of MP trees found by dnapars, by swapping parsimony-optimal substructures between them^30^. Next, GCtree ranks the resulting collection of maximally parsimonious trees. For our analysis we configured the ranking to first minimize the number of mutations reverting to the naïve ancestral state, then optimize a branching process likelihood, then a context-based Poisson likelihood, and finally minimize the number of unique sequences in the trees to arbitrarily break any remaining ties. The branching process likelihood uses observed abundances of genotypes to implement the intuition that sequences observed with higher abundance should have more mutant offspring. Trees which follow this intuition have greater branching process likelihood^25^. The context-based Poisson likelihood measures how inferred mutations in the tree agree with context-dependent mutation rates and targeting probabilities of an S5F model^23,30^. Dnapars infers ancestral states, marking sites for which multiple ancestral states are possible under maximum parsimony. GCtree attempts to enumerate all possible maximally parsimonious ancestral states for ranking. If this is not possible for a topology found by dnapars, a single maximally parsimonious ancestral reconstruction is chosen.

#### GC selection metrics

NDS was calculated by splitting each GC into root-clades (branches that stem directly from the unmutated ancestor) then dividing the number of cells in the largest lineage by the total sequenced cells in that GC. REI is calculated as follows: For each node X in a phylogeny, the REI represents the sum of the number of descendants of node X weighted according to their mutational distance from node X using a decay factor 𝜏 = 0.5, such that cells at 0, 1, 2, … nucleotide distance from node X are weighted 1, 0.5, 0.25, …; the sum of weighted descendants is then divided by the total number of cells in the GC. A phylogeny in which all cells have the same sequence therefore has an REI of 1.0. This metric relies on the assumption is that GC B cells cease SHM when undergoing rapid clonal burst-type expansion^42,96^. The NDS and REI scores are computed on these trees using utilities that are part of our computational pipeline stored on GitHub (https://github.com/matsengrp/gcreplay/blob/main/analysis/NDS-LB.ipynb).

#### Calculation of intrinsic mutability

BCR libraries were processed utilizing the pRESTO suite tools^97^. We adhered to the Illumina MiSeq 2x250 BCR mRNA pipeline. Initially, low-quality reads were filtered out, followed by primer masking and unique molecular identifier (UMI) quantification to correct sequencing errors and PCR amplification biases. Sequences were paired, and consensus sequences were constructed for R1 and R2 reads. These consensus sequences were further paired and assembled into contiguous sequences. Finally, identical sequences were collapsed and quantified. Sequences represented by at least two reads were used for downstream analysis.

BLAST was used to identify sequences with full-length matches for the regions around the CRISPR/Cas9-induced indels that defined the passenger alleles using a 90% identity threshold. The corresponding reads were then subject to a series of filters: the identifying sequence could only be found once in the read, the read could only have one additional indel, and the read could have at most 9 mutations and at most 9 N positions. As is typical in the field^23,98^ the SHM model was described in terms of a “substitution probability,” namely the probability of the new base conditioned on there being a substitution, and a per-site mutability estimate of having a mutation at each position.

In order to combine information across multiple runs for each of the heavy and light chains, we used a Poisson modeling strategy that allowed for the read depth and the overall mutation load to differ between experiments, as well as the mutation rate to differ between sites (further details are provided in https://github.com/matsengrp/gcreplay/blob/main/passenger/igh_passenger_aggregate.ipynb). The per-site rates are parameterized using softmax, resulting in a collection of rates across the sites that sum to 1. A small number of *Igk* positions that had > 0.001% Ns (*Igk* nucleotides 295, 306, 307, 319, 321), indicative of potential sequencing errors, were excluded and replaced by the value obtained using from the five-mer model^23^.

#### Affinity fitness landscape modelling

The minimal mathematical model for the distribution of affinities over time was developed as follows: we denote the log_10_ affinity change with respect to naive as 𝑥 and specify an evolution equation for the probability density of affinities 𝑝(𝑥, 𝑡) at each time 𝑡. There are two key parameter functions that influence the time evolution of 𝑝(𝑥, 𝑡) in our model. First, a function 𝑞(𝑥, 𝑦) represents the mutational flux (mutation rate per unit 𝑥 and 𝑦) from affinity state 𝑥 to affinity state 𝑦, and can be specified (up to a multiplicative scale) by weighting the distribution of log_10_-affinity effects measured in DMS by the mutabilities of each mutation measured from the passenger mouse experiment. Second, an *affinity fitness landscape* 𝑓(𝑥) specifies the intrinsic fitness for a cell with affinity 𝑥, and is an unknown function of key interest, as it can be used to predict GC competition between cells of different affinities. The interpretation of this fitness is competitive, so that the Malthusian growth rate of a cell with affinity 𝑥 at time 𝑡 is given by 𝑓(𝑥) − 𝑓̅(𝑡), where 𝑓̅(𝑡) is the mean fitness in the population at time 𝑡. Because the affinity distribution evolves through time, 𝑓̅(𝑡) is expected to increase with time, so that a cell with a given affinity becomes less competitive as its competitors tend to improve in affinity. Assuming a large population, the deterministic evolution equation is

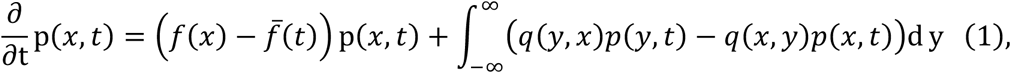

where the first term models birth and death, and the second term models mutation into and out of affinity state 𝑥^45^. Because the replay mouse has monoclonal naïve cells starting GCs, we have the initial condition 𝑝(𝑥, 𝑡_0_) = δ(𝑥), i.e. a Dirac mass at initial time 𝑡_0_. Note that this equation is nonlinear due to the dependence of the mean fitness 𝑓̅ on 𝑝, via 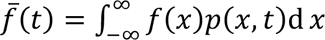. Also note that this is an equilibrium population size model, which can be seen by integrating both sides of the evolution equation (1) over 𝑥, which shows the time derivative of the normalization of 𝑝 vanishes identically.

We now detail how we solve the evolution equation, fitting it to the time-course experiment data at time points 5, 8, 11, 14, 17, 20, and 70 dpi. We implement a custom Crank-Nicholson numerical PDE solver as a differentiable program^99^ and use maximum likelihood estimation (MLE) to infer the parameters of the evolution equation that best fit the time-course data, chiefly the fitness landscape 𝑓(𝑥) which we represent nonparametrically as a smooth monotonic function. The time-course data 𝒟 consists of a set of sampling times 𝑡 and a set of real-valued affinities 𝒳 at each such time. The sample size of a sample (𝑡, 𝒳) ∈ 𝒟 is |𝒳|. The log-likelihood of 𝑓(𝑥) given the time-course data is

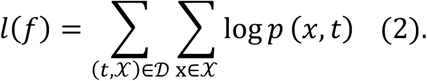

This likelihood represents the likelihood of observing the time course data given a density of affinities through time 𝑝(𝑥, 𝑡). This 𝑝(𝑥, 𝑡), in turn, is derived deterministically by solving the PDE (1): although 𝑓does not appear explicitly on the right-hand side of (2), 𝑝 is a functional of 𝑓. We perform MLE by backpropagating through the PDE solver to maximize the log-likelihood, using an entropic penalty to encourage the fitted mutational flux profile *q* to match the empirical data from passenger mouse and DMS, and a spline penalty on *f* to suppress high-frequency artifacts that aren’t supported by the data fit. More details are available in https://github.com/matsengrp/gcreplay/blob/main/analysis/affinity-fitness-response.ipynb.

#### Stochastic extinction calculation for survivorship bias effects

We now derive a master equation approach for the stochastic extinction probability 𝑟,(𝑥, 𝑡) of a lineage with affinity state 𝑥 at time 𝑡 before sampling time 𝑇. This lineage undergoes a continuous-state birth-death-mutation process^100^ according to birth rate λ(𝑥) = f(𝑥) – f_0_, death rate µ(𝑡) = f̅(𝑡) – f_0_, and mutation flux q(𝑥, 𝑦) to affinity state 𝑦. We define reference fitness 𝑓_0_ corresponding to millimolar affinity. Note that the expected growth rate is f(𝑥) − f̅(𝑡), matching the previous deterministic model. Whence the forward master equation is

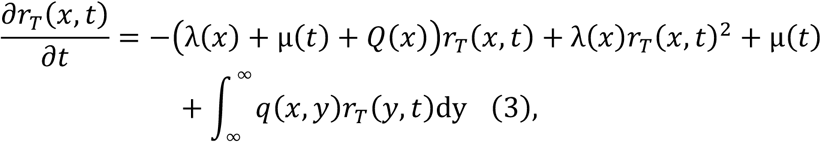

where 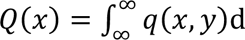 y is the total mutation intensity. Such a master equation can be derived by considering the probability of events (birth, death, mutation, or nothing) that can happen in a small time interval Δ𝑡 (such that at most one event occurs), then evaluating r.(x, t + Δt) and taking Δ𝑡 to be infinitesimal, yielding a partial differential equation. The first term corresponds to nothing happening; it is linear in the extinction probability because the lineage remains non-extinct after the infinitesimal time interval. The second term corresponds to a birth event in the infinitesimal time interval; it is quadratic in the extinction probability because both daughter lineages must independently go extinct for the parent lineage to go extinct. The third term corresponds to a death event, given simply as the instantaneous Poisson death rate because the lineage is then extinct. The integral in the last term describes a mutation event to any possible other affinity state; it is linear in the extinction probability at the alternative affinity state because the mutated lineage would then have to go extinct. A full derivation for a related problem can be found in Kuhnert et al. 2016^101^ and a briefer derivation for a birth-death process can be found in Barido-Sottani et al. 2020^102^. We solve the equation backward in time from final condition 𝑟,(𝑥, 𝑇) = 0, again using our custom Crank-Nicholson solver.

Equipped with a numerical solution to this master equation, we compute the ancestral affinity distribution (the distribution conditioned to leave descendants at the sample time 𝑇) by weighting 𝑝(𝑥, 𝑡) by the probability of survival to time 𝑇 and renormalizing

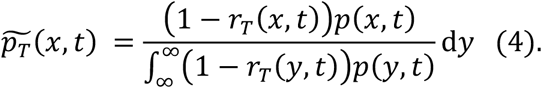

We used this forward master equation approach to compute the probability 𝑟_r_(𝑥, 𝑡) of stochastic extinction before sampling time 𝑇 = 15 dpi or 𝑇 = 20 dpi.

#### Bayesian phylogenetic analysis

We performed Bayesian phylogenetic analysis using BEAST v1.10.4. Nucleotide sequences were analyzed under an HKY substitution model with a strict molecular clock and a constant coalescent demographic model. Markov chain Monte Carlo (MCMC) sampling was conducted for 25 million iterations, with trees sampled every 10,000 steps. After discarding the first 96% of samples as burn-in, the remaining 100 trees were used for the push-of-the-past analysis. The XML template file for the BEAST analysis can be found at https://github.com/matsengrp/gcreplay/blob/main/data/beast/beast_templates/constantsize_histlog.template.patch

#### Other software

Flow cytometry data were analyzed using FlowJo v. 10. Plots and statistical analyses not included in Jupyter notebooks were generated using Graphpad Prism v. 10. Figures were edited for appearance using Adobe Illustrator.

#### Declaration of generative AI and AI-assisted technologies in the manuscript preparation process

During the preparation of this work the author(s) used Claude and ChatGPT as aids in coding and text editing (but not to generate text). After using this tool/service, the author(s) reviewed and edited the content as needed and take(s) full responsibility for the content of the published article.

## SUPPLEMENTAL INFORMATION

**Figure S1.**
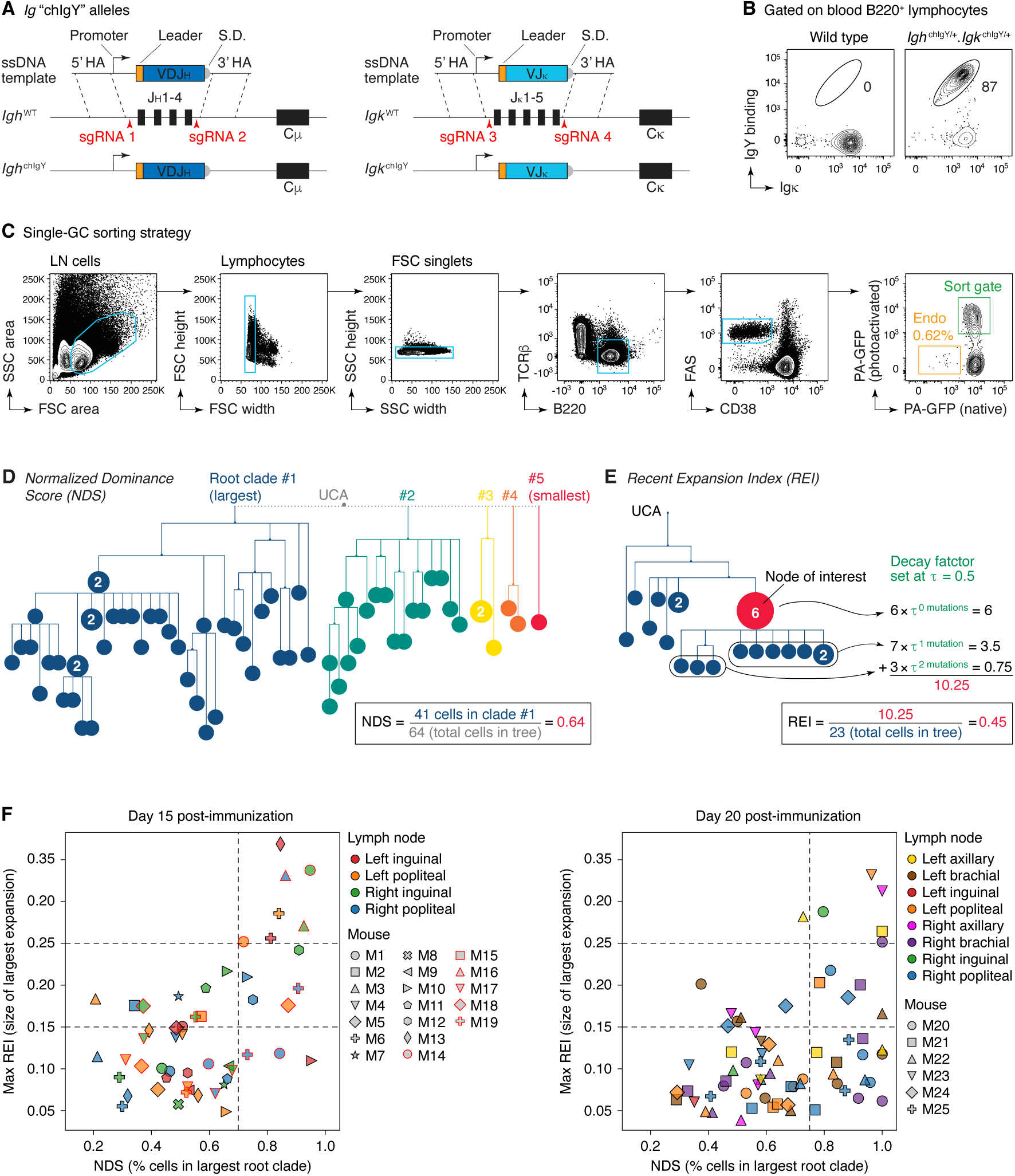
Parallel replay of germinal center evolution. **(A)** Design of the *Igh* and *Igk* “chIgY” alleles. CRISPR/Cas9 genome targeting with single-stranded DNA templates was used to replace the endogenous JH and Jκ segments with pre-rearranged V(D)J genes. HA, homology arm; S.D., splice donor. **(B)** Flow cytometry of blood B cells from wild-type and chIgY mice, showing expression of the rearranged V(D)J genes (inferred from the ability of B cells to bind IgY). **(C)** Sorting strategy for the parallel replay experiment. FACS plots show a representative LN fragment. Endo, residual endogenous GC B cells derived from the CD23-Cre.*Bcl6*^flox/flox^ host. **(D)** Schematic representation of the normalized dominance score (NDS) calculation. NDS is equal to the percentage of all cells in a GC that belong to the largest root clade (dark blue). The size of the smaller clades is not included in the calculation. **(E)** Schematic representation of the recent expansion index (REI) calculation. For each node X in a phylogeny, the REI represents the sum of the number of descendants of node X weighted according to their mutational distance from node X using a decay factor 𝜏 = 0.5, such that cells at 0, 1, 2, … nucleotide distance from node X are weighted 1, 0.5, 0.25, …; the sum of weighted descendants is then divided by the total number of cells in the GC. A phylogeny in which all cells have the same sequence therefore has an REI of 1.0. **(F)** Distribution of NDS and REI scores for GCs from 15 and 20 dpi as in Fig. 1E,F, but colored by mouse and LN of origin; each symbol represents one GC.

**Figure S2.**
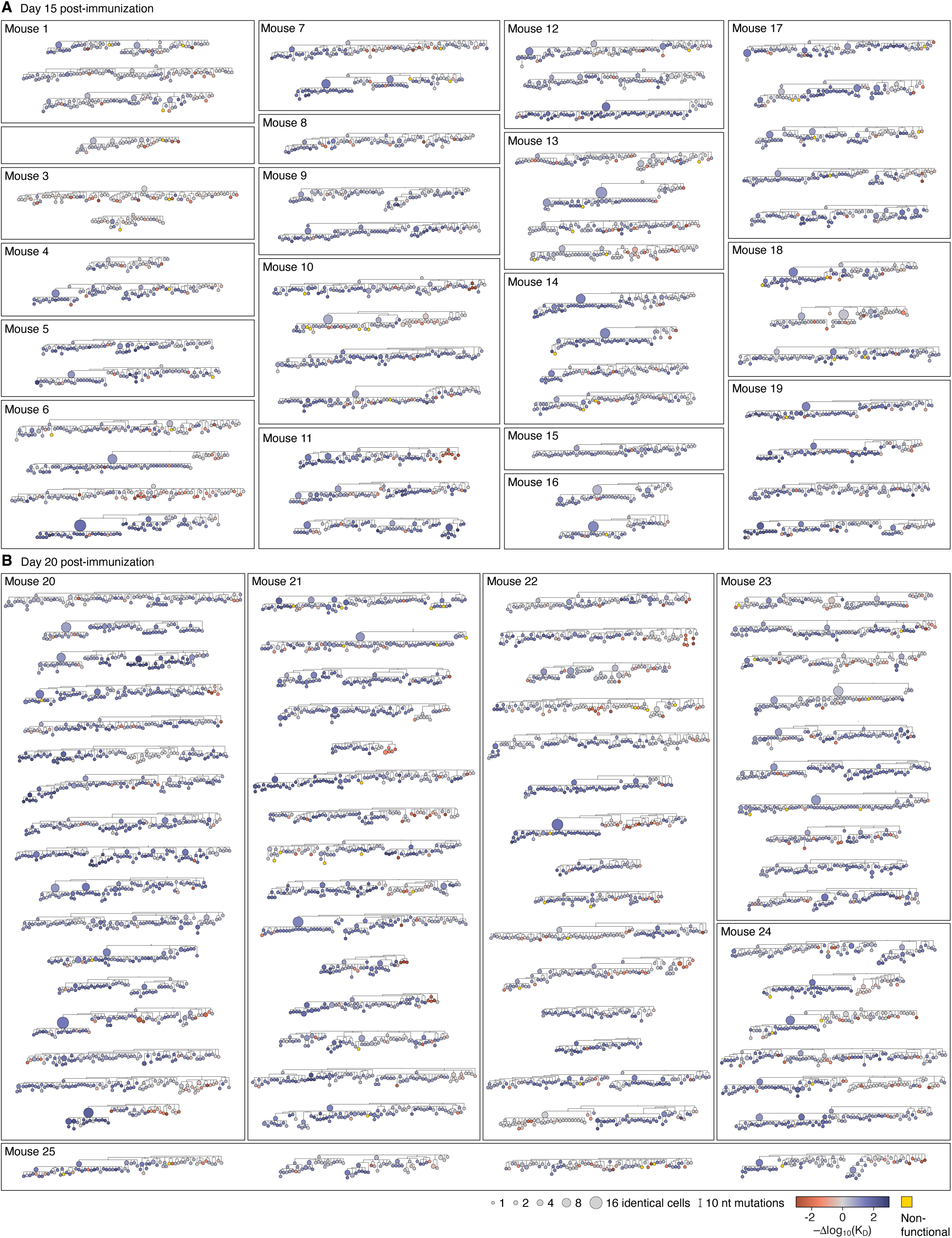
Individual germinal center phylogenies. Diagrams of all 119 phylogenetic trees inferred from the *Igh+Igk* sequences obtained from each photoactivated GC at **(A)** 15 and **(B)** 20 days post-immunization. Boxes indicate the mouse from which each GC was sorted. Nodes are colored by their relative affinity from the naive precursor (–Δlog10(KD)), see Fig. 2 for details.

**Figure S3.**
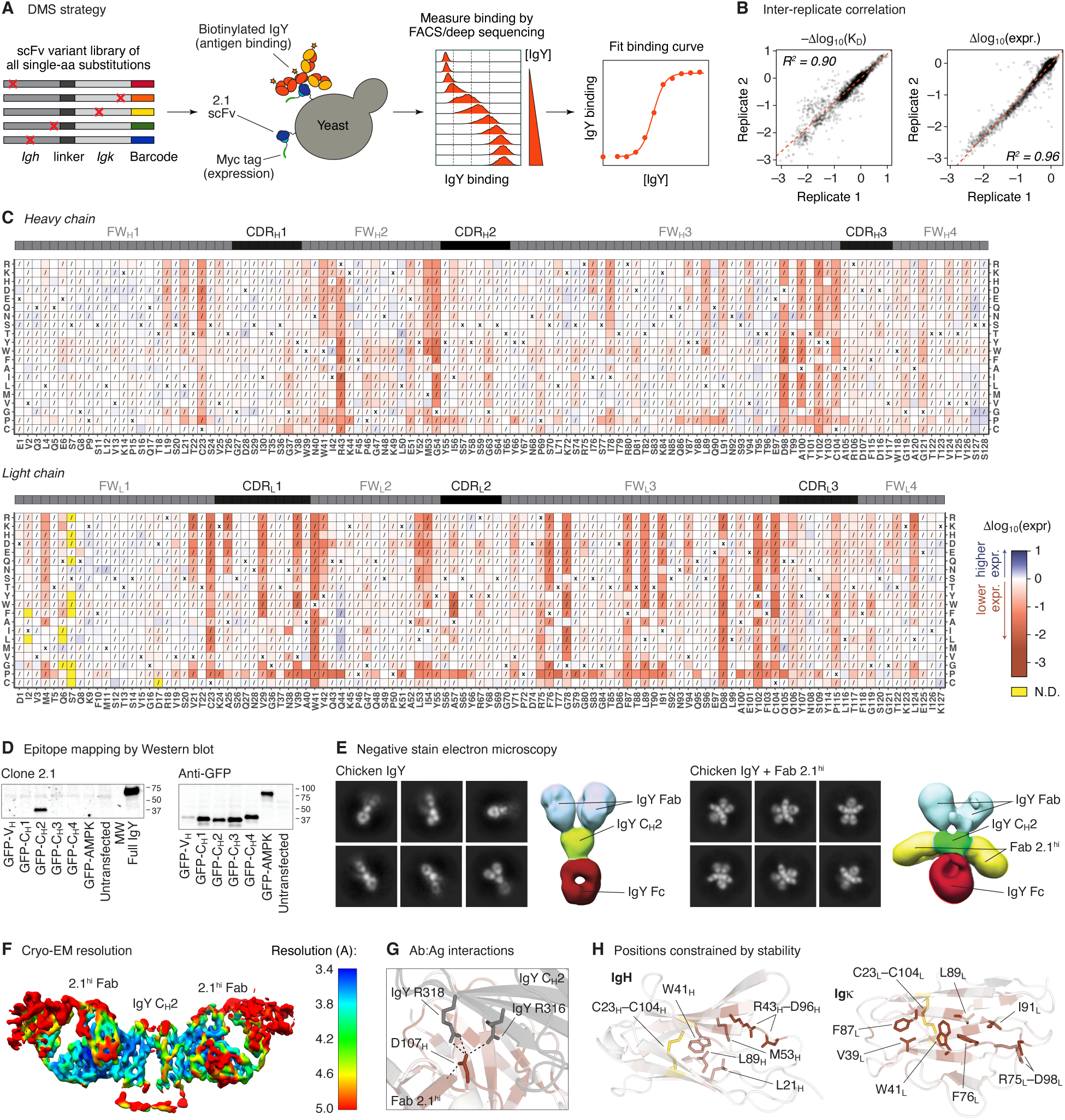
Deep mutational scanning and structure of clone 2.1. **(A)** Experimental setup for the DMS experiment. **(B)** Correlation in mutation effects on CGG binding affinity (left) and scFv expression level (right) from independently generated and assayed mutant libraries. **(C)** Heatmap showing the effects of individual amino-acid replacements on surface expression of 2.1 scFv by yeast display. Each square represents a different replacement. Squares with asterisks indicate the original amino acid in clone 2.1. Squares marked with “X” indicate the original amino acid in clone 2.1. Squares with slashes indicate amino acid replacements distant more than one nucleotide mutation from the naïve sequence. Yellow squares were not detected in the DMS experiment. Upper bar shows Kabat CDR and framework (FW) regions in black and gray, respectively. An interactive version of this heatmap is available at https://matsengrp.github.io/gcreplay/interactive-figures/mutation-heatmaps/naive_reversions_first.html. **(D)** Coarse mapping of the epitope of clone 2.1. 293T cells were transfected with constructs encoding each Ig domain of IgYH (or AMPK as a control) fused to GFP. Cell extracts were probed by Western blot with recombinant clone 2.1 mAb (left) or with polyclonal anti-GFP to detect expression of the construct (right). MW, molecular weight ladder, not visible by Western blot. **(E)** Negative-stain 2D classes and 3D reconstructions (colored by domain/subunit) of unliganded chicken IgY (left) and the 2.1^hi^ Fab:IgY complex (right) confirming Fab binding to CH2. **(F)** Cryo-EM reconstruction of the 2.1^hi^ Fab:IgY complex colored by local resolution following IgY CH2 and 2.1^hi^Fab local refinement. **(G)** Key inferred electrostatic interaction between clone 2.1 and IgY. Backbone and side chains colored by mean ΔKD for all replacements at each position. Color scale as in Fig. 2A. Potential salt bridge interactions are shown as dotted lines. Mutation of D107H to anything other than an acidic amino acid (D107_H_E) results in at least 1 log_10_ decrease in binding affinity. **(H)** Mapping of selected residues that strongly affect antibody surface expression when mutated. Backbone and side chains colored by mean Δ expression for all replacements at each position. Color scale as in (B); disulfide bridges shown in yellow.

**Figure S4.**
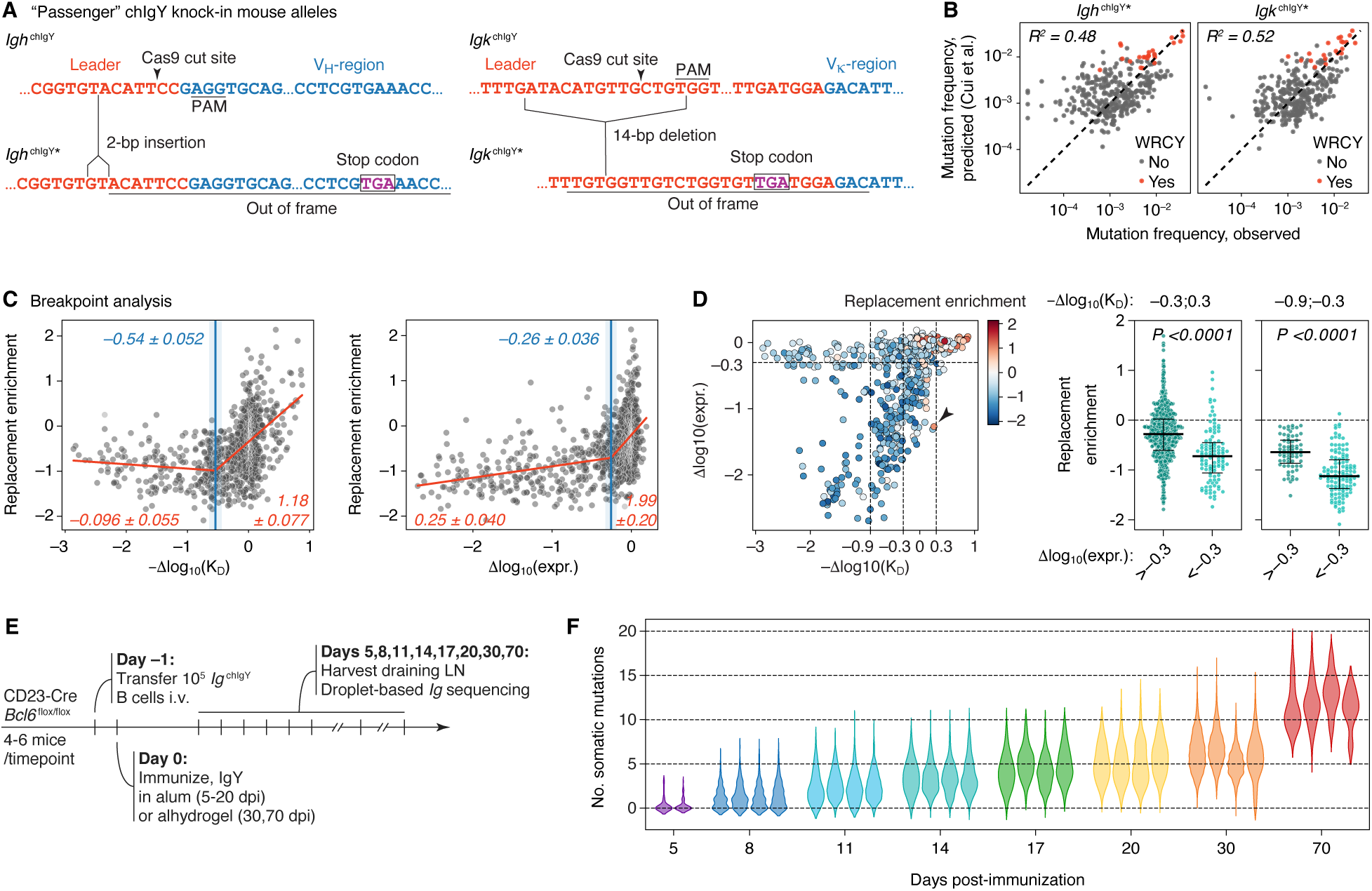
Selection of individual amino acid replacements. **(A)** Sequences of the “passenger” *Igh*^chIgY^* and *Igk*^chIgY^* alleles generated by CRISPR/Cas9-mediated cleavage on the *Ig*^chIgY^ background. **(B)** Comparison of relative mutation frequencies observed in passenger allele mice *in vivo* with predictions made using the five-mer model^26^. Each symbol represents one nucleotide position of the respective *Ig* sequence. C/G pairs within R<u>G</u>YW AID hotspot motifs are highlighted in red. **(C)** Segmented regression analysis using the piecewise-regression package in Python. A two-segment fit was chosen by model comparison, and the results of that fit are shown here. **(D)** Each replacement, plotted in terms of its effect on affinity and expression, and colored according to replacement enrichment. Arrowhead indicates an exceptional replacement that is enriched even though it leads to a marked loss of expression. **(E)** Layout of the time-course experiment. **(F)** Distribution of somatic mutations at the indicated time points post-immunization. Each violin represents one mouse. Data for 5-20 dpi were obtained together in a single experiment; data for 70 dpi is from a separate experiment. Two mice (of a total of 9) were excluded from the 70 dpi violin plots for insufficient cell yield; cells from these mice were included in the bulk analysis.

**Figure S5.**
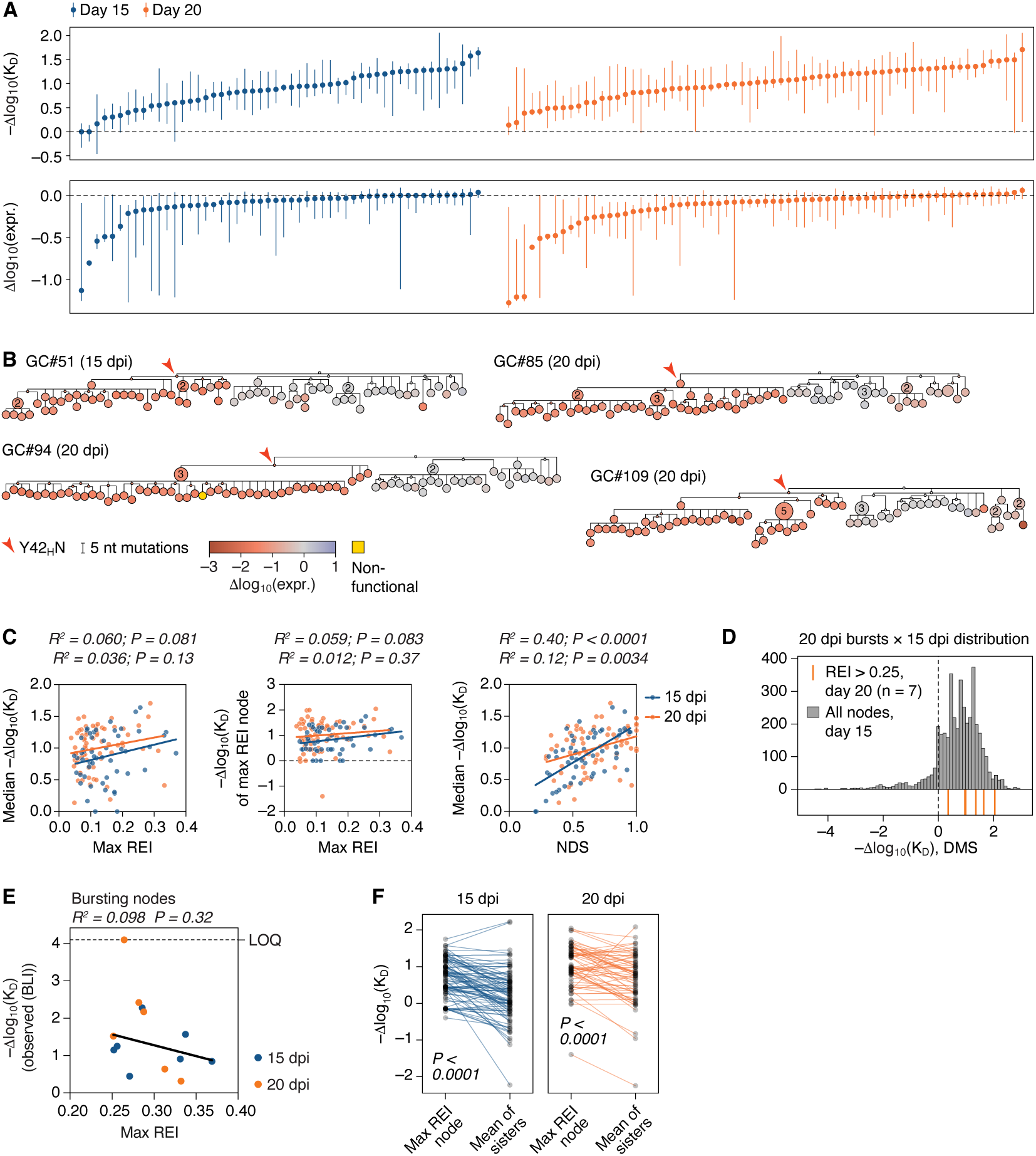
Drivers of affinity maturation and maintenance of Ig expression. **(A)** Median (± range) of Δaffinity (top) and Δexpression (bottom) for all GCs in the replay experiment. Each symbol represents one GC. **(B)** Examples of GC phylogenies carrying the expression-impairing Y423HN replacement. Trees are colored by Δexpression. **(C)** Correlations between phylogenetic selection parameters for each GC (max REI and NDS) and median Δaffinity or the Δaffinity corresponding to the sequence of the max REI node). Each symbol represents one GC. Trend lines are for each time point. R^2^ and P-values are for Pearson correlation and are given for 15 dpi (top row) and 20 dpi (bottom row). **(D)** Overlay of the distribution of Δaffinities for clonal burst nodes obtained at 20 dpi (REI >0.25; orange lines) compared to the distribution of Δaffinities for all nodes (observed and inferred) from 15 dpi (grey bars). **(E)** Correlation between BLI-measured Δaffinities and the Max REI in the GC (corresponding to the REI of the sequence of the bursting B cell itself) of each of the bursting nodes in Fig. 5D. Solid black line is the linear trend. LOQ, limit of quantitation, when Fab off-rate is too long to be determined. R^2^ and slope p-value calculations exclude the Fab with Δaffinity >LOQ. **(F)** Statistical comparison of the affinity of the max REI node in each replay GC and the mean affinity of its “sister” nodes, as defined in Fig. 5G. P-values are for the Wilcoxon signed-rank test.

**Figure S6.**
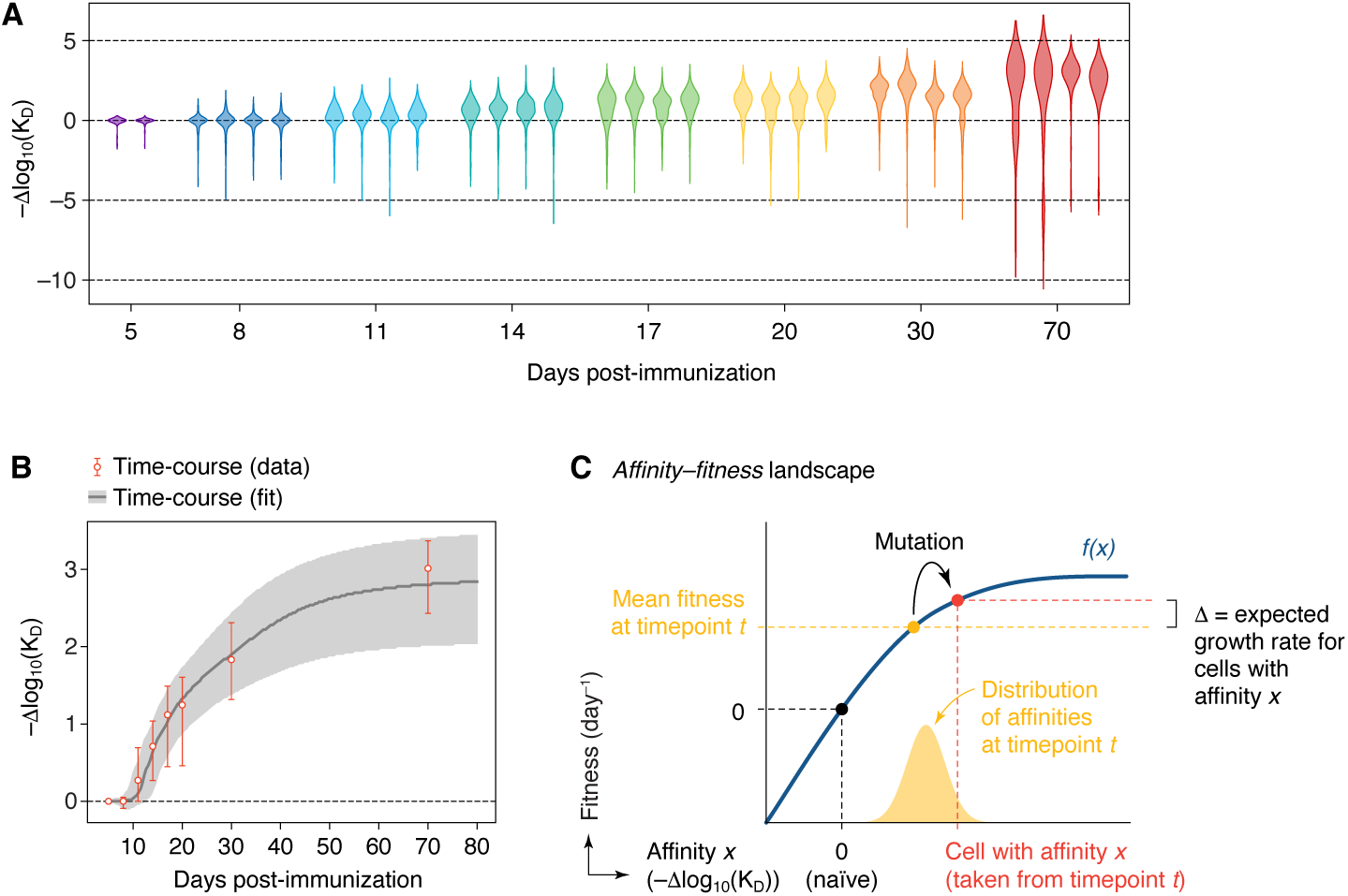
Progression of affinity over time in bulk-sorted GC B cells. **(A)** Left, distribution of DMS-inferred affinities at the indicated time points post-immunization. Each violin represents one mouse. Data for 5-20 dpi were obtained together in a single experiment; data for 70 dpi is from a separate experiment. Two mice (of a total of 9) were excluded from the 70 dpi violin plots for insufficient cell yield; cells from these mice were included in the bulk analysis. Right, distribution of DMS-inferred affinities at 70 dpi, after exclusion of cells below that persisted in GCs despite having lost all detectable binding to IgY (given that these cells can potentially be explained by co-option of GC lineages to bind other antigens^106,107^ in a host mouse strain unable to generate GCs of its own). For each mouse, the upper mode of the distribution was identified, and a threshold was placed manually (red lines in the panel on the right), below which cells were excluded from further analysis. **(B)** Given the consistency of affinity distributions across individual mice, we aggregated cells from each time point to approximate a single longitudinal sequence of affinity distributions sampled throughout the time-course. Graph shows median and IQR of this distribution overlaid on the prediction generated by the fitness landscape model. **(C)** Schematic representation of the affinity–fitness landscape, adapted from Held et al.^47^ This landscape translates changes in affinity (X-axis) into changes in “evolutionary fitness” (Y-axis), corresponding roughly to the expansion rate expected for a population of B cells with affinity X when compared to a naïve competitor population with affinity = 0, in the absence of further mutation. Operationally, the difference (Δ) between the fitness of a population of B cells with affinity *x* sampled at time-point *t* (red) and the mean fitness of all cells at time point (orange) corresponds to the expected exponential growth rate (or decay rate, when negative) for that population over time. The detailed shape of the function *f(x)* specifies how fitness responds to marginal affinity increases and how this responsiveness attenuates as one moves up in the affinity axis. Mathematical details of the model are elaborated in the Methods.

**Supplemental Table 1.** Data collection, processing, model refinement and validation.

| **Map** | **IgY + Clone 2.1 Fab**<br>**(local refinement)** |
| --- | --- |
| EMDB | EMD-70353 |
| **Data collection** |  |
| Microscope | Thermo Fisher Talos Arctica |
| Voltage (kV) | 200 |
| Detector | Gatan K2 Summit |
| Recording mode | Counting |
| Nominal magnification | 36,000x |
| Movie micrograph pixelsize (Å) | 1.15 |
| Dose rate (e<sup>-</sup>/[(camera pixel)*s]) | 6.95 |
| Number of frames per movie micrograph | 47 |
| Frame exposure time (ms) | 200 |
| Movie micrograph exposure time (s) | 9.5 |
| Total dose (e<sup>-</sup>/Å<sup>2</sup>) | 50 |
| Defocus range (µm) | -0.8 to -2.5 |
| **EM data processing** |  |
| Number of movie micrographs | 10,007 |
| Number of molecular projection images in map | 346,259 |
| Symmetry | C1 |
| Map resolution (FSC 0.143; Å) | 4.0 |
| Map sharpening B-factor (Å<sup>2</sup>) | -171 |
| **Structure building and validation** |  |
| Number of atoms in deposited model |  |
| IgY CH2 | 1,434 |
| Clone 2.1 Fab | 3,400 |
| glycans | 28 |
| MolProbity score | 1.22 |
| Clashscore | 1.36 |
| Map correlation coefficient | 0.73 |
| EMRinger score | 1.63 |
| d FSC model (0.5; Å) | 4.3 |
| RMSD from ideal |  |
| Bond length (Å) | 0.007 |
| Bond angles (°) | 1.203 |
| Ramachandran plot |  |
| Favored (%) | 94.96 |
| Allowed (%) | 5.04 |
| Outliers (%) | 0.00 |
| Side chain rotamer outliers (%) | 0.00 |
| Cβ outliers (%) | 0.00 |
| PDB | TBD |

**Supplemental Spreadsheet 1 (online). Biolayer interferometry results.**

**Supplemental Spreadsheet 2 (online). CRISPR guide and insert sequences used to generate *Ig*^chIgY^ mice.**

## Notes

https://matsen.group/gcreplay/key-files/#manuscript-figures

### Summary of Updates

Major changes were made to the structure of the text describing the mathematical modeling (Fig. 6). Minor changes were made elsewhere.

